# A meta-analysis of single visit pollination effectiveness

**DOI:** 10.1101/2021.03.12.432378

**Authors:** M.L. Page, C.C. Nicholson, R.M. Brennan, A.T. Britzman, J. Greer, J. Hemberger, H. Kahl, U. Müller, Y. Peng, N.M. Rosenberger, C. Stuligross, L. Wang, L.H. Yang, N.M. Williams

## Abstract

Many animals provide essential ecosystem services in the form of plant pollination. A rich literature documents considerable variation in the single visit pollination effectiveness of different plant visitors, but this literature has yet to be comprehensively synthesized. We conducted a hierarchical meta-analysis of 193 studies and extracted 1716 single visit effectiveness (SVE) comparisons for 252 plant species. We paired SVE data with visitation frequency data for 75 of these studies. Given the global dominance of honeybees in pollinator communities, we used these data to ask: 1) Do honeybees (*Apis mellifera*) and other floral visitors vary in their SVE?; 2) To what extent do plant and pollinator attributes predict the difference in SVE between honeybees and other visitors?; and 3) Is there a correlation between floral visitation frequency and SVE? We found that honeybees were significantly less effective than the most effective non-honeybee pollinator. Although not significantly different, honeybees also tended to be less effective than the mean community effectiveness. Honeybees were less effective as pollinators of crop plants and when compared to birds and other bees. Visitation frequency and pollination effectiveness were positively correlated, but this trend was largely driven by data from communities where honeybees were absent, suggesting that honeybees generally combine high visitation frequency and lower SVE. Our study demonstrates that non-honeybee floral visitors are highly effective pollinators of many crop and non-crop plants. While the high visitation frequency typically displayed by honeybees undoubtably makes them important pollinators, we show that honeybees are slightly less effective than the average pollinator and rarely the most effective pollinator of the plants they visit. As such, honeybees may be imperfect substitutes for the loss of wild pollinators and safeguarding global crop production will benefit from conservation of non-honeybee taxa.

**Open Research Statement:** Although we are fully committed to data transparency, we are also aware of different research teams working on related meta-analyses. As such, we prefer to wait until our paper is accepted to make data publicly available but are happy to share data upon request. Data will be permanently archived on Figshare following acceptance.

## Introduction

Over 70% of plants depend to some degree on animal pollinators to successfully reproduce (Ollerton et al. 2011). Among the diversity of pollinators, there is large variation in the ways different taxa contribute to pollination. Biodiversity ecosystem-function theories posit that diverse communities will optimize niche space, allowing the entire community to be more productive (Loreau et al. 2001, Cardinale et al. 2012), but a few dominant species can provide the majority of ecosystem functioning (Lohbeck et al. 2016). For pollination, the functional contribution of different pollinators is measured by the quality (single visit effectiveness, SVE) and quantity (frequency) of their visits to plant reproductive success (King et al. 2013).

Pollination effectiveness describes the per-visit contribution of floral visitors to plant pollination (Inouye et al. 1994). A long history of studies within the botanical and evolutionary ecology literature documents wide variation in single visit effectiveness (SVE) between plant visitors (e.g., Herrera 1987, King et al. 2013, Page et al. 2019). To some extent, variation in pollination effectiveness reflects the wide range of methods used to measure SVE (Ne’eman et al. 2010), such as single visit pollen deposition (King et al. 2013), the number of developed pollen tubes within styles (Zhang et al. 2015), and/or fruit or seed set (Vicens and Bosch 2000). However, evidence for variation in SVE comes from numerous individual studies focused on the reproduction of single plant species in specific contexts. Meta-analysis could broaden our understanding of whether and why particular pollinators are more effective than others and help evaluate persistent hypotheses about the factors that influence pollination.

In particular, the high visitation frequency typical of honeybees (*Apis mellifera*) has been hypothesized to drive their functional contribution as pollinators (Hung et al. 2018), regardless of their per-visit effectiveness. Indeed, an extensive literature on pollinator importance – the product of per-visit effectiveness and relative visitation rates of different pollinators (King et al. 2013, Ballantyne et al. 2015) – concluded that pollinators that visit more frequently are generally more important (Vásquez et al. 2012). This conclusion suggests that numerical dominance generally outweighs among-species variation in SVE, but it does not indicate that more frequent visitors are more effective on a per-visit basis.

However, the processes governing pollination effectiveness and visitation frequencies may not be independent of one another. A few key studies have suggested that a pollinator’s SVE may be positively correlated with its visitation frequency (e.g., Vásquez et al. 2012, Ballantyne et al. 2017). Ballantyne et al. (2017) found a positive correlation between a pollinator’s visit frequency and pollination effectiveness when comparing 23 plant species, likely because bees were both highly effective and highly frequent visitors compared to other floral visitors. Positive correlations between pollination effectiveness and visit frequency could also occur if pollinators who visit frequently do so to the exclusion of other plant species. For example, high floral constancy could minimize heterospecific pollen transfer resulting in more effective pollination (Morales and Traveset 2008). On the other hand, high visitation rates may be the result of many quick and ineffective visits (Ohara et al. 1994) and contribute a negative or non-significant effect on reproductive success in many contexts (e.g., Sáez et al. 2014, reviewed in Willcox et al. 2017). In the case of honeybees, apiary management and location may drive high visitation rates relative to other pollinators. As such, correlations between visitation rates and pollination effectiveness could be obscured by the presence of managed honeybees.

Despite their high visitation frequencies, the effectiveness of honeybees relative to other pollinators remains unclear. Outside of their native range, honeybees lack the evolutionary history with endemic plants that could have selected for increased pollinator effectiveness (Javorek et al. 2002). Because honeybees are floral generalists that visit a high proportion of available plants in ecosystems across the globe (Hung et al. 2018), they may not be particularly effective at pollinating specific flowering species. For example, honeybees sometimes ‘rob’ plants (Irwin et al. 2010) and efficiently extract and groom pollen from plants without depositing the pollen they extract (Westerkamp 1991) or collect nectar without contacting reproductive structures (Vicens and Bosch 2000). However, honeybees can be highly effective pollinators even for plants with which they have no shared evolutionary history (Wist and Davis 2013), suggesting that shared evolutionary history is not a prerequisite to effective pollination.

Understanding pollinator effectiveness has important practical implications for safeguarding the production of pollinator-dependent crops. Highly effective non-honeybee pollinators are important for ensuring crop pollination in the face of global change (Rader et al. 2013) and functionally diverse pollinator communities can increase crop pollination (Woodcock et al. 2019). Furthermore, pollination dynamics may differ in cultivated settings because interspecific plant competition, the spatial arrangement of flowers, and the pollinator taxa that provide pollination may vary across agricultural and natural landscapes (Harrison et al. 2018).

We used a meta-analysis of the pollination effectiveness literature to address three key questions. First, how does the SVE of honeybees compare to that of other floral visitors? We hypothesized that honeybees would exhibit lower SVE relative to other pollinators because honeybees are broad generalists and might lack the evolutionary history with specific plants that could tune pollination effectiveness. Second, to what extent do plant and pollinator attributes predict the comparative SVE of honeybees? Specifically, we evaluated the role of broad taxonomic pollinator groups (e.g., bees, birds, etc.), crop status (crop vs. non-crop plant species), and whether plant species exist within the native range of honeybees. We hypothesized that the SVE of honeybees would be lower compared to other bees in particular, in crop systems, and for plant species outside the native range of honeybees. Third, is there a correlation between floral visitation frequency and SVE? We evaluated this question separately for communities where honeybees were present or absent, hypothesizing a positive correlation between visitation frequency and SVE that would be reduced in contexts where honeybees dominate. Although previous studies have synthesized subsets of the pollination effectiveness literature (notably, Hung et al. 2018, Földesi et al. 2020), this paper is the first meta-analysis to comprehensively synthetize published results concerning single visit effectiveness.

## Methods

### Study Screening

We performed a Web of Science (WoS) search using a multiterm query (Appendix S1: Fig. S1) designed to capture the highly variable terminology describing pollination effectiveness detailed in Ne’eman et al. (2010). In May 2020, this search yielded 1,036 results. One of us (MP) screened the abstracts found by WoS to determine whether they potentially contained single visit effectiveness (SVE) data. This yielded 388 papers. We also performed a Google Scholar search of the literature using a similar multi-term query (Appendix S1: Fig. S1), which yielded 116 additional papers. We found 62 papers from the reference sections of previously included papers. After removing duplicates and reading abstracts, we identified 468 papers which seemed appropriate for a more thorough screening.

We followed the PRISMA protocol for collecting and screening data from the literature (Appendix S1: Fig. S1; Moher 2009). To be included in our analysis, the paper had to contain empirical data on the per-visit contribution of at least one free-foraging visitor to plant reproduction. We considered pollen deposition, percent fruit set, fruit weight, and/or seed set as measures of SVE. Most studies were conducted with intact flowers, but we also included data from experiments that used the “interview stick” method (in which a cut flower was presented to potential visitors). We did not include estimates of SVE based on equations or model outputs nor did we include data from trials that manipulated dead bees to deposit pollen. We extracted means, sample sizes, and measures of error (e.g., standard deviation, standard error) directly from the text of the paper or from graphs using WebPlotDigitizer (v. 4.4, Rohatgi 2020). When lower and upper error estimates were not symmetrical, we used the upper error estimate. When possible, we converted measures of error to standard deviation. When a paper did not report sample sizes, error, or other important information, we contacted the study authors. If we were unable to retrieve or estimate information on mean effectiveness and error, we excluded the paper from our analysis. After screening papers, 193 studies remained in our analytical dataset. We also extracted data on study year and location, plant species, plant family, whether the plant species was a crop-plant, pollinator taxon, pollinator group (e.g., bird, fly, bee), and the native range of pollinator and plant species. We determined range status to biogeographical realms by looking up the nativity of each taxon in the scientific literature and using occurrence records on GBIF. If papers reported SVE outcomes from multiple sites or years, we extracted these data as separate outcomes and dealt with their non-independence statistically (see below).

We collected information on the visitation rates of pollinators if it was reported for the same plant species for which pollinator effectiveness data were reported. This rate could be reported as the number of visits to a focal flower or patch of flowers per unit time or the number of flowers visited per unit time and/or per unit area. We did not include data on the relative abundance of different visitors, unless data were collected in a homogeneous landscape (like an orchard) in which most visitors would have been visiting the focal plant species. If a study reported visitation data, we matched that data to the corresponding SVE data from the same study and plant species. Perfect matches required that pollinator taxa were reported to the same taxonomic resolution and that data were collected in the same year and location. When more than one measure of visit frequency was reported we preferentially used data on the number of visits to a focal flower per unit time. When more than one measure of SVE was reported, we preferentially chose whichever measure was better represented in our data, such that pollen deposition data were chosen over seed set data and seed set data were chosen over fruit set data.

### Meta-analysis

To address questions about the single visit effectiveness of honeybees and non-honeybees, we defined the effect size as the standardized mean difference (SMD, i.e., Hedges *g* (Hedges 1981)) of SVE values between honeybee and non-honeybees for each unique study, plant, site, and year combination. We chose to use Hedge’s g over other effect sizes because it is commonly used in the ecology literature for comparing two means (Nakagawa and Santos 2012), and it includes a correction for small sample sizes, which occurred with our data. Following Hung et al. (2018), we calculated effect sizes for two separate comparisons: (1) the difference between honeybees versus the most effective non-honeybee taxon and (2) the average difference between honeybees and non-honeybee taxa (hence, ‘average effectiveness’). The SMD value is > 0 when other pollinators are more effective than honeybees and < 0 if the opposite occurs. We calculated each effect size in *R* (R Core Development Team 2020) using the *escalc* function in the ‘*metafor’* package (v. 2.1-0, Viechtbauer 2010).

We fit meta-analytic and meta-regression multilevel linear mixed-effects models, using the *rma.mv* function in the ‘*metafor’* package (v. 2.1-0, Viechtbauer 2010). We used three random effects to control for non-independence of effect sizes collected from the same study or plant species: study ID, plant species, and an observation level ID. We used phylogenetic comparative methods (Cornwell and Nakagawa 2017) to account for non-independence that may arise due to shared evolutionary history of focal plants by including a phylogenetic covariance matrix. The phylogeny used to compute a phylogenetic covariance matrix (Zanne et al. 2014) was constructed using the package ‘*brrranching’* (v. 0.6.0, Chamberlain 2020), and branch lengths were set following Grafen’s method (Grafen 1989) using the *R* package ‘*ape’* (v. 5.1, Paradis and Schliep 2019)). Despite slightly higher AIC values and larger P values (Appendix S2: Table S1), we present results from models including phylogenetic controls to fully account for non-independence due to shared ancestry (Chamberlain et al. 2012). With this mixed-effects structure, we specified four models, which include an intercept only model (i.e., overall meta-analytic model), and three meta-regression models for different fixed effects/moderators: (1) pollinator taxonomic group, (2) whether the plant was a crop plant (crop status), and (3) for native plants, whether it was in the honeybee’s native range (range status). We follow Hung et al. (2018) and define the West Palearctic as the honeybees’ native range (Ruttner 1988).

To test whether there was a relationship between a pollinator taxon’s single visit effectiveness and visit frequency, we calculated Pearson’s correlation coefficients (r) for the relationship between visit frequency and pollinator effectiveness for each unique study, plant, site, and year combination in which there were at least five pollinator taxa represented. After calculating correlation coefficients, we used the *escalc* function in the *metafor* package to calculate Fisher’s r-to-Z transformed correlations and corresponding sampling variances. Using the same multilevel linear mixed-effects model structure and phylogenetic controls as described above we generated three models. The first model was an intercept-only model to test for the overall relationship between a pollinator’s visit frequency and single visit effectiveness. The second model compared three categories against one another: studies where honeybees were present, studies where honeybees were absent, and studies where we artificially removed all points corresponding to honeybees (re-calculating effect sizes as detailed above). We generated this third category to determine whether the patterns we observed were solely driven by honeybees themselves or whether there might also be indirect effects of honeybee presence on the relationship between visit frequency and single visit effectiveness. The third model tested whether there was an interaction between crop status and honeybee presence.

### Tests for publication bias

Publication bias was assessed based on visual inspection of funnel plots for each model (Appendix S1: Fig. S4, S5) and via a modified Egger’s test (Egger et al. 1997, Sterne and Egger 2005) on meta-analytic residuals in which effect size precision (sqrt(1/variance)) is included as a moderator (Nakagawa and Santos 2012). A significant slope for precision would indicate statistically significant funnel asymmetry after controlling for all other variables in the model. We considered analyses to be biased if the intercept differed from zero at P = 0.10 (as in Egger et al. 1997).

## Results

We built a dataset of 1716 SVE records (i.e., average effectiveness values for pollinators visiting plants) drawn from 193 peer-reviewed and published studies (Appendix S1: Table S1). Research was conducted on every ice-free continent, with most work occurring in the Nearctic (N = 68) or West Palearctic (N = 42) over a period of 39 years, from 1981 to 2020 (Fig. 1). Many studies (34) investigated pollination of more than one plant species (range: 2-23), with a total of 252 plant species assessed belonging to 67 families. Across plants and studies, relative effectiveness values were normally distributed (Appendix S1: Fig. S3a), but most pollinators (53%) were less effective than the mean effectiveness of all visitors, compared to 44% who were more effective than the mean. For studies that reported visit frequency data (N = 75), the distribution of relative visit frequency values was skewed to the right (Appendix S1: Fig. S3b), such that only 27% of visitors visited more frequently than the mean visit frequency. Within studies that reported paired effectiveness and visit frequency data, honeybees were the most frequent visitor 28% of the time but the most effective pollinator only 9% of the time.

**Figure 1.**
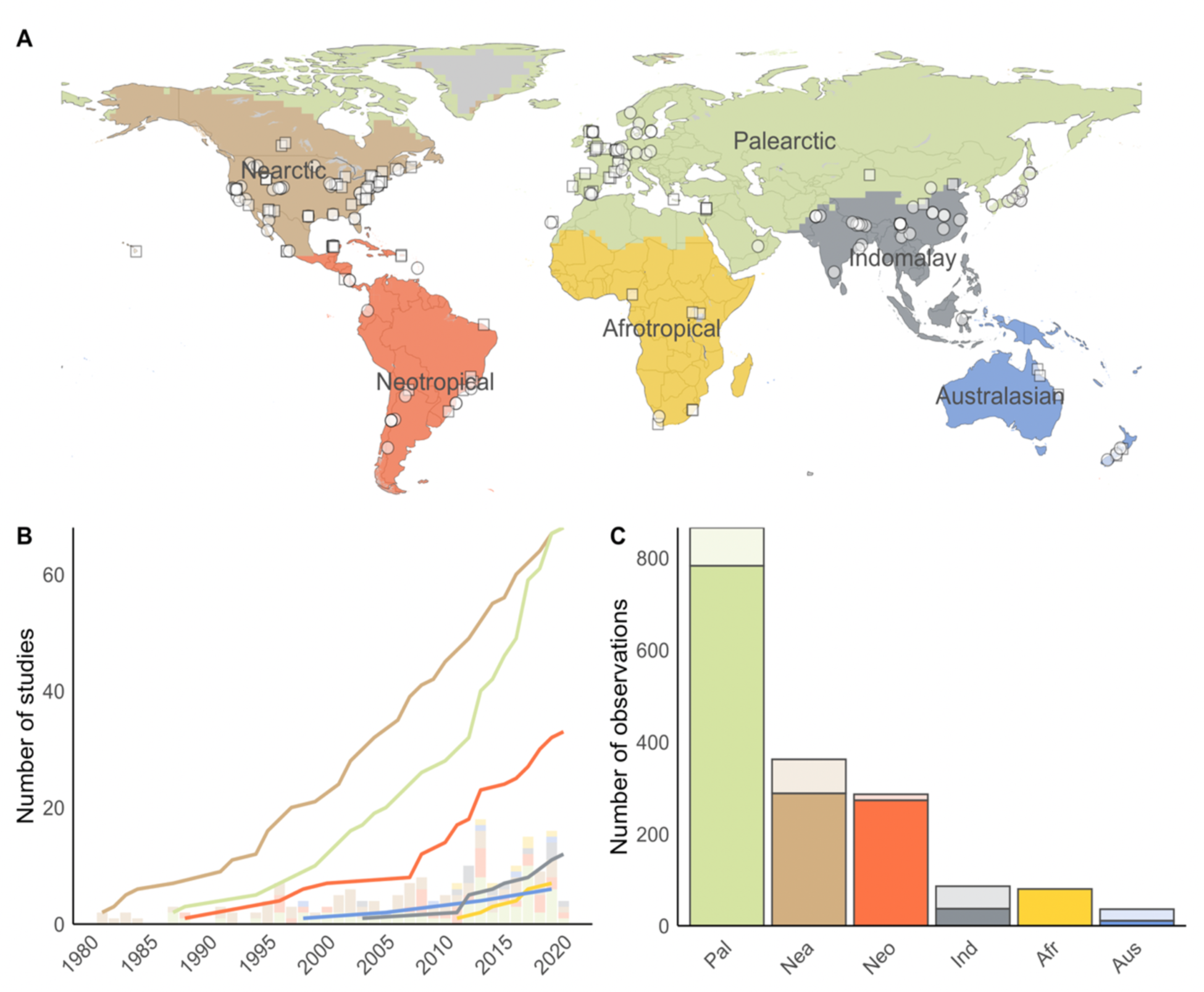
The research into single visit effectiveness (SVE) is geographically widespread and has progressed consistently over time. (A) Map of study locations depicting whether research recorded SVE measures for honeybees and other taxa (squares) or if honeybees were the sole taxon or absent (circles). (B) Trends in SVE research show the cumulative number of studies per region (lines) and the annual number of studies (rug). (C) Some studies have more than one SVE observation (e.g., multiple pollinators visiting multiple plants); observation totals varied across regions and based on whether plants were native (dark colors) or non-native (lighter colors).

### How does the SVE of honeybees compare with other floral visitors?

A total of 84 studies reported comparisons between *A. mellifera* and at least one other taxon. These studies focused on 96 plant species and include crops (N = 32) and non-native plant species (N = 22) (Appendix S1: Fig. S2). From these comparative studies, 621 individual effect sizes were obtained and summarized for each combination of plant and pollinator group within a study. This yielded 186 effect sizes comparing the most effective non-honeybee pollinator and honeybees (Most Effective Pollinator (MEP) comparisons) and 186 effect sizes comparing the average effectiveness of all non-honeybee pollinators and honeybees (Average Effective Pollinator (AEP) comparisons). When comparing overall study-level effect sizes, we found that non-honeybee pollinators were more effective than honeybees. This outcome was statistically significant for Most Effective Pollinator comparisons (Appendix S2: Table S1; overall standardized mean difference (SMD): 0.497, [0.211, 0.783 95% CI]; P = 0.001), but not Average Effective Pollinator comparisons (SMD: 0.207, [-0.094, 0.508]; P = 0.177). The data showed little evidence of publication bias in terms of funnel plot asymmetry of meta-analytical residuals as revealed by plot inspection (Appendix S1: Fig. S4). Results from Egger’s tests suggested little to no degree of asymmetry for our overall meta-analytic model (MEP: P = 0.06; AEP: P > 0.10).

### To what extent do plant and pollinator attributes predict the comparative SVE of honeybees?

Computing effects separately for each pollinator group revealed that the type of pollinator moderated the comparative SVE of honeybees (Fig. 2). The most effective bees and birds were significantly more effective than honeybees (Fig. 2a; bee SMD: 0.665, [0.463, 0.868]; P < 0.001 & bird SMD: 2.269, [1.457, 3.082]; P < 0.001). For average effectiveness comparisons, only birds were significantly more effective than honeybees (Fig. 2b; SMD: 1.344, [0.667, 2.020]; P < 0.001). Although the average non-*Apis* bee tended to be more effective than honeybees, this difference was not statistically significant (SMD: 0.247, [-0.094, 0.588]; P = 0.156). However, significant differences were detected in models without phylogenetic controls (Appendix S2: Table S1; SMD: 0.322, [0.137, 0.506]; P = 0.001), suggesting that additional data might confirm this trend. At the study level, 61% of effect sizes compared other bees and honeybees; we therefore focus subsequent analyses on bees.

**Figure 2.**
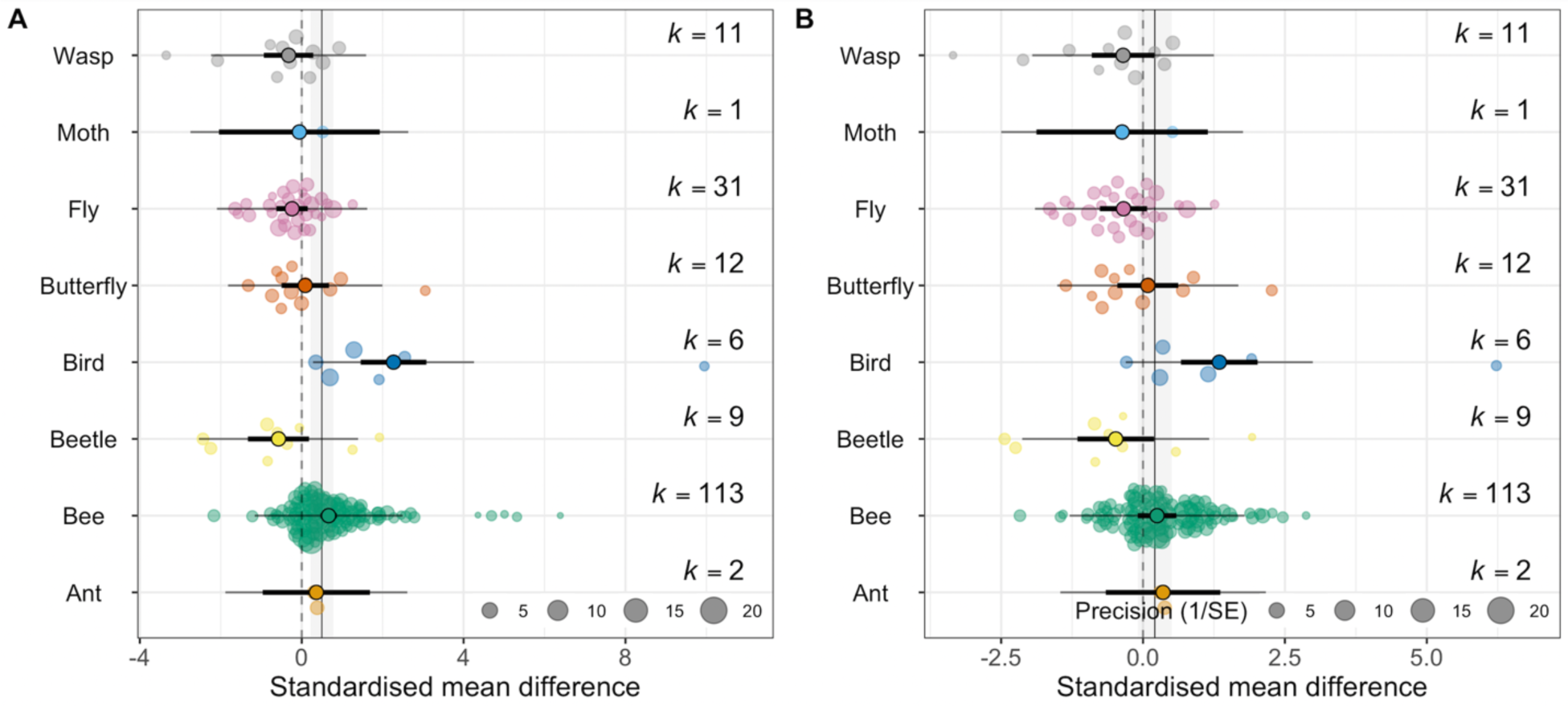
Meta-regression results for single visit effectiveness differences between honeybees and different pollinator taxa. Effect sizes (Standard mean difference, SMD) were calculated (A) between honeybees and the most effective non-honeybee taxon within each group and (B) between the average effectiveness across all non-honeybee taxa within each group for a given plant-study. Meta-analytic means are represented as point estimates with their 95% CI (thick lines) and prediction intervals (thin lines). Point estimates from meta-regressions are depicted with their 95% CI (thick lines) and prediction intervals (thin lines). Individual effect sizes are scaled by their precision (1/SE). Positive SMD values (points to the right of zero) indicate that other pollinators were more effective than honeybees.

The most-effective bees were more effective pollinators of crops than honeybees (Fig. 3a; SMD: 0.786, [0.328, 1.244]; P = 0.001); this was true for average effectiveness comparisons as well (Fig. 3b; SMD: 0.511, [0.137, 0.886]; P = 0.007). For non-crop plants, honeybees were marginally less effective than the most effective other bees (Fig. 3a; SMD: 0.413, [-0.033, 0.859]; P = 0.069), but were not significantly different than the average bee pollinator. The most-effective bees were better pollinators of native plants than honeybees (Fig. 4a); this was true for plants occurring within (SMD: 0.608, [0.021, 1.195]; P = 0.042) and outside (SMD: 0.683, [0.105, 1.261]; P = 0.021) *Apis mellifera*’s native region (West Palearctic). Honeybees were comparable to the average SVE of bees (Fig. 4b), inside and outside their native range.

**Figure 3.**
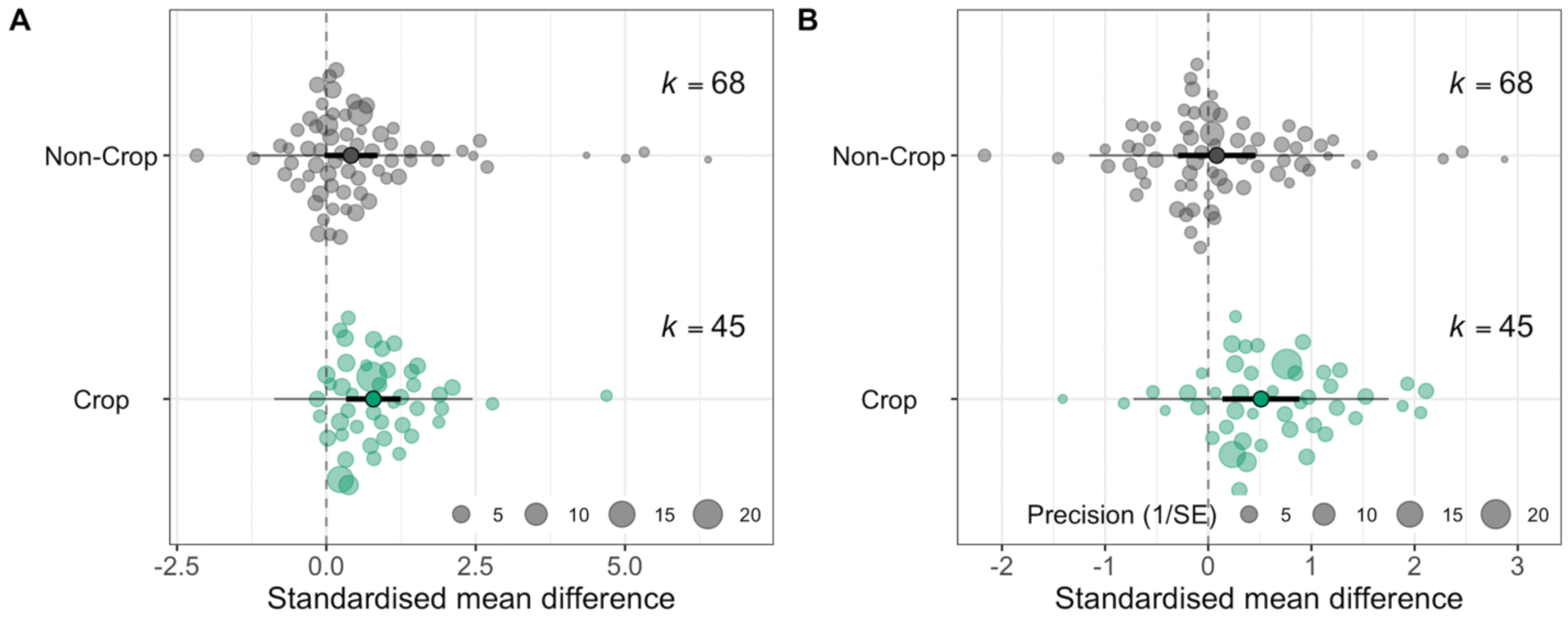
Meta-regression results for crop single visit effectiveness differences between honeybees and non-honeybee bees. Effect sizes (Standard mean difference, SMD) were calculated (A) between honeybees and the most effective non-honeybee bee and (B) between the average effectiveness across all non-honeybee bees for a given plant-study. Effect sizes were compared for non-crop (gray circles) and crop species (green circles). Meta-analytic means are represented as point estimates with their 95% CI (thick lines) and prediction intervals (thin lines). Individual effect sizes are scaled by their precision (1/SE). Positive SMD values (points to the right of zero) indicate that other bees were more effective than honeybees.

**Figure 4.**
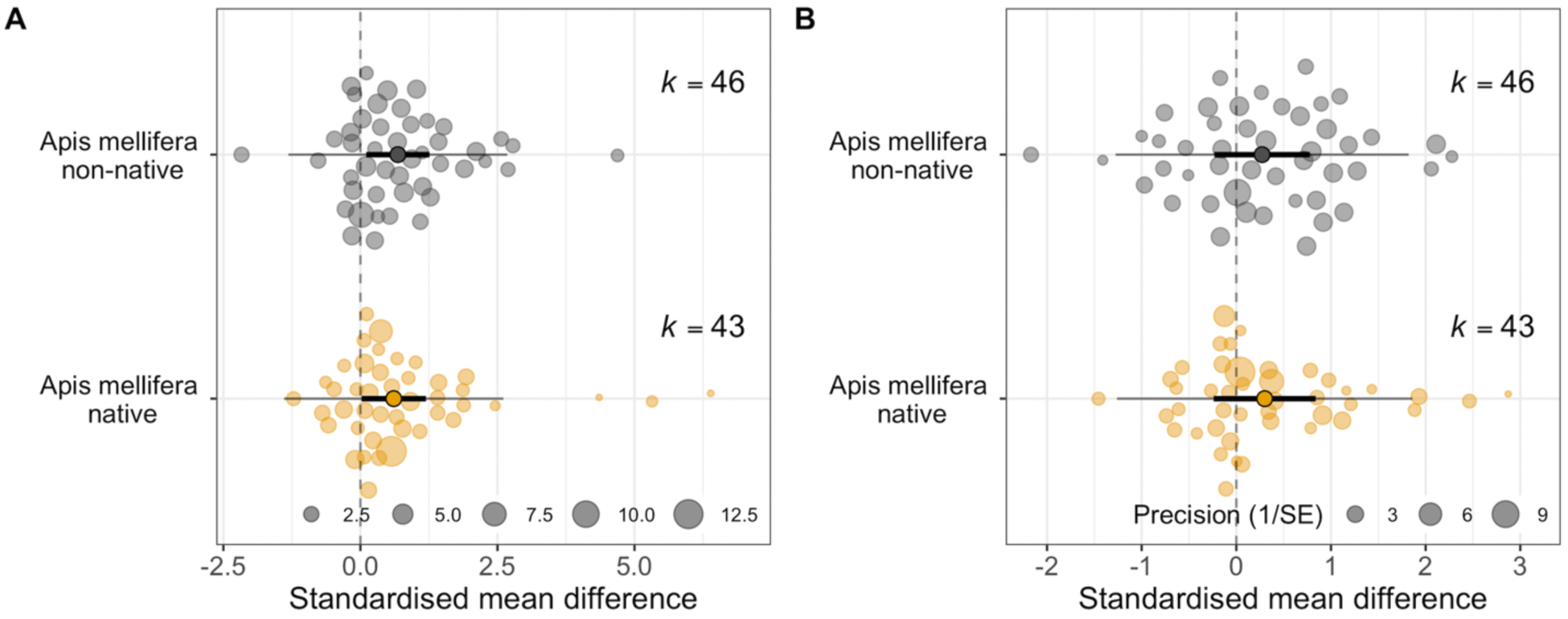
Meta-regression results for native plant single visit effectiveness differences between honeybees and non-honeybee bees. Effect sizes (Standard mean difference; SMD) were calculated (A) between honeybees and the most effective non-honeybee bee and (B) between the average effectiveness across all non-honeybee bees for a given plant-study. Effect sizes were compared outside (gray circles) and inside (orange circles) the honeybee native range. Meta-analytic means are represented as point estimates with their 95% CI (thick lines) and prediction intervals (thin lines). Individual effect sizes are scaled by their precision (1/SE). Positive SMD values (points to the right of zero) indicate that other bees were more effective than honeybees.

### Is there a correlation between floral visitation frequency and SVE?

Overall, there is a positive relationship between visit frequency and single visit effectiveness (Estimate: 0.600 [0.127, 1.074 95% CI]; P = 0.013). However, data from systems in which honeybees are absent drive this positive result. When honeybees are present, there is no relationship between visit frequency and effectiveness (Fig. 5; Estimate: 0.390 [-0.131, 0.911]; P > 0.05) and this lack of a significant relationship persisted when we artificially removed data corresponding to honeybee visits. We observed a positive association between visit frequency and SVE only when *Apis mellifera* was actually absent (Fig. 5; Estimate: 0.758 [0.242, 1.273]; P = 0.004). There was no interaction between honeybee presence and crop status (Appendix S1: Fig. S6). An Egger’s test confirmed minimal publication bias (P > 0.10). There were a few outliers to the right of funnel plots (Appendix S1: Fig. S5). Removing these data did not influence our findings.

**Figure 5.**
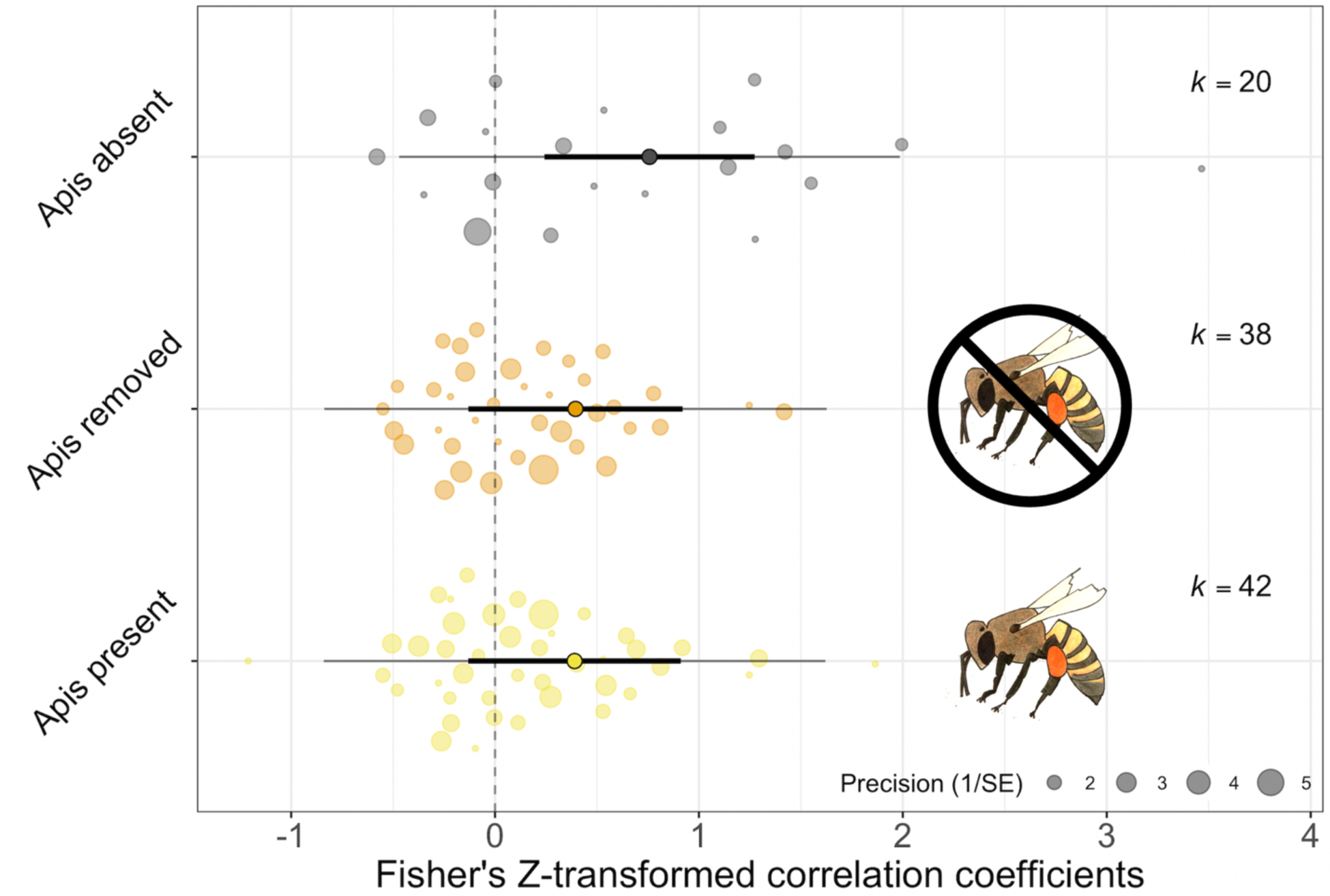
Meta-regression results for the relationship between a pollinator’s visit frequency and single visit effectiveness for studies with and without honeybees present. Effect sizes (Fisher’s Z-transformed correlation coefficients) were compared for systems where honeybees were absent (gray circles), systems where honeybees were present (yellow circles, also indicated by honeybee icons), and systems where honeybees were present when data were collected, but we artificially removed data corresponding to their visits and re-calculated correlation coefficients (orange circles, also indicated by crossed-out honeybee icons). Estimates are shown with their 95% CI (thick lines) and prediction intervals (thin lines). Effect sizes are scaled by their precision (1/SE).

## Discussion

Our meta-analysis supports the hypothesis that honeybees are frequently not the most effective pollinator of plants globally. Across six continents and hundreds of plant species, honeybees showed significantly lower single visit effectiveness than the most effective pollinator in a community (Fig. 2). Although not statistically different, honeybees also tended to be less effective than the community mean. This general pattern is likely driven by comparison of honeybees against birds and other bees. The most effective bird and bee pollinators were significantly more effective than honeybees, as were the average bird pollinators. Although not statistically different, honeybees were also less effective than the average non-*Apis* bee. The finding that birds are more effective than honeybees is based on only six studies that were likely focused on flowers frequently pollinated by birds. Nevertheless, it supports the idea that plants adapted to bird pollination have traits that enhance pollination by birds at the expense of pollination by bees (Castellanos et al. 2006). Although data for non-bee taxa were relatively sparce, honeybees were equally as effective as the average and most effective ant, beetle, butterfly, fly, moth, and wasp pollinators, confirming that non-bee insects can be important pollinators (Rader et al. 2020). Our results bolster initial work summarizing honeybee pollination effectiveness (Hung et al. 2018) and demonstrate that honeybees are less effective than many other visitors and at best average.

Analysis of crop plants also revealed important differences between honeybees and non-*Apis* pollinators. Despite their abundance in commercial cropping systems, honeybees are less effective crop pollinators than the most effective bee pollinators and the average non-honeybee bees (Fig. 3). This finding supports the idea that the importance of honeybees as crop pollinators derives largely from their numerical dominance as crop visitors (Hung et al. 2018). Our analysis adds robust evidence to a growing consensus that wild bees have the potential to contribute greatly to agricultural pollination. Indeed, wild bee species richness, functional diversity, and visit rates increase crop yield (Blitzer et al. 2016, Woodcock et al. 2019), and the use of managed honeybee hives might not compensate for losses in wild bee species richness and abundance (Mallinger and Gratton 2015, Pérez-Méndez et al. 2020). As such, managed honeybees alone may be insufficient to meet the increased pollination demands of global agricultural production (Aizen and Harder 2009) and our results validate the importance of actions to promote resilient native bee communities within agricultural lands (Isaacs et al. 2017).

Honeybees were consistently less effective compared to other bees when pollinating native plants both inside and outside the honeybee’s native range (Fig. 4). This result is not entirely surprising based on what we know about the co-evolution of plants and pollinators. The non-honeybee bee community may contain specialists sympatric with their host plants. Meanwhile, if honeybees are broad generalists, selective pressure might be less consistent, even within the native range of honeybees. Furthermore, if the morphological features relevant to pollination are relatively consistent across plants within the same genus or family, insects may be capable of pollinating novel plant species. For example, *Prunus* spp. occur in Europe and North America and *Osmia* spp. are highly effective pollinators of *Prunus* tree crops in both regions (Vicens and Bosch 2000, Bosch et al. 2006), despite the fact that North American *Osmia* spp. do not have shared evolutionary history with the *Prunus* species introduced as tree crops.

We found an overall positive relationship between visit frequency and single visit pollinator effectiveness, but this relationship was largely driven by data from systems in which honeybees were absent (Fig. 5). The overall positive correlation suggests that more frequent visitors are also more effective, but this result should not be interpreted to indicate that visitation frequency is an adequate proxy for overall pollination importance (Vásquez et al. 2012, Ballantyne et al. 2017). This positive correlation may suggest that pollinators which visit frequently do so to the exclusion of other plant species, such that they display high floral constancy. High floral constancy may indicate that visitors gather and transport more conspecific pollen (Brosi and Briggs 2013). Although the pollen loads of visitors do not always adequately predict effective pollination (Adler and Irwin 2006), high conspecific pollen transport likely predisposes visitors to higher pollination effectiveness on average.

The finding that honeybees erode this otherwise positive correlation suggests that this hyper-generalist species is often a numerically dominant visitor with modest effectiveness and may modify the pollination context for plant communities. Interestingly, when comparing systems with and without honeybees, visit frequency and pollination effectiveness do not positively correlate even when we artificially remove the data on honeybees and re-calculate correlation coefficients. This result suggests that honeybee presence may indirectly influence the relationship between visitation frequency and pollination effectiveness by altering the visitation patterns and effectiveness of other plant visitors. High honeybee visitation frequencies may indicate that honeybees efficiently extract nectar and pollen without also efficiently depositing the pollen they extract (Westerkamp 1991, Wilson and Thomson 1991, Goodell and Thomson 1997). If honeybees deplete floral nectar, this could make plants less attractive to other common visitors (Hansen et al. 2002) and alter their visit behavior and effectiveness (Thomson 1986). If they extract large amounts of pollen (Cane and Tepedino 2017), this could reduce the amount available for collection and deposition by other pollinators (Harder and Barrett 1995).

There are several potential limitations of our study and possibilities for future work. First, we only included measures of female reproductive success in assessing pollination effectiveness (e.g., pollen deposition, seed set). The proportion of extracted pollen that is successfully transferred to stigmas may be a better assessment of the overall reproductive contribution of different taxa (Parker et al. 2016), because pollen that is removed but not successfully transferred represents a loss to male fitness (Harder and Thomson 1989, Minnaar et al. 2019). Unfortunately, data on such transfer dynamics are much rarer in the literature. Second, there are likely other factors about plant and pollinator taxa that moderate the effects we observe but which we do not test in this study, for example, functional traits such as plant and pollinator specialism. We hope our study will motivate other researchers to pair our data with trait databases and information on single visit pollen removal to further investigate the factors that influence effective pollination.

As honeybees become increasingly dominant globally, the abundance and species richness of other pollinators visiting plants is expected to decrease (Valido et al. 2019). If honeybees replace visits from other pollinators, whether through competitive displacement or otherwise (Herrera 2020), these changes in community composition may have cascading effects on plant pollination, reproduction, and persistence (Gómez et al. 2010). Species loss and fluctuations in the abundance of important pollinators can imperil ecosystem service delivery (Cardinale et al. 2012, Winfree et al. 2015). Even rare species are important to ecosystem functioning (Winfree et al. 2018) and functionally diverse pollinator assemblages enhance plant community diversity (Fontaine et al. 2005). If honeybees are not particularly effective, it will be key to understand whether and how honeybees influence the visitation frequencies and effectiveness of other pollinators. Another key question is whether honeybees can compensate for the inferior quality of their visits with increased visit frequency, which can occur (Sun et al. 2013). Ultimately, some plants will thrive as their visitor community becomes increasingly dominated by honeybees, while others may experience declines. Given increasing honeybee dominance, it will be important to identify and protect diverse and effective pollinator communities especially when confronted with ineffective substitutes.

## Acknowledgements

We would like to thank Priya Shukla who helped with data collection and provided thoughtful feedback during a graduate seminar in which this meta-analysis was first proposed. The quality of this manuscript benefited greatly thanks to comments from Elizabeth Crone and anonymous reviewers. MLP and CCN are joint first authors. All authors contributed to data collection, idea generation, and manuscript revisions. MLP and CCN wrote the manuscript and analyzed the data. MLP was supported by a U.S. Department of Defense National Defense Science and Engineering Graduate (NDSEG) Fellowship. CCN was supported by a USDA-ARS Non-Assistance Cooperative Agreement No. 58-2030-8-031 awarded to NMW. JG, HK, and CS were supported by National Science Foundation Graduate Research Fellowships. NMW was partially funded by NSF DEB 1556885. We declare no conflict of interest.

## Appendix S1

**Table S1.**
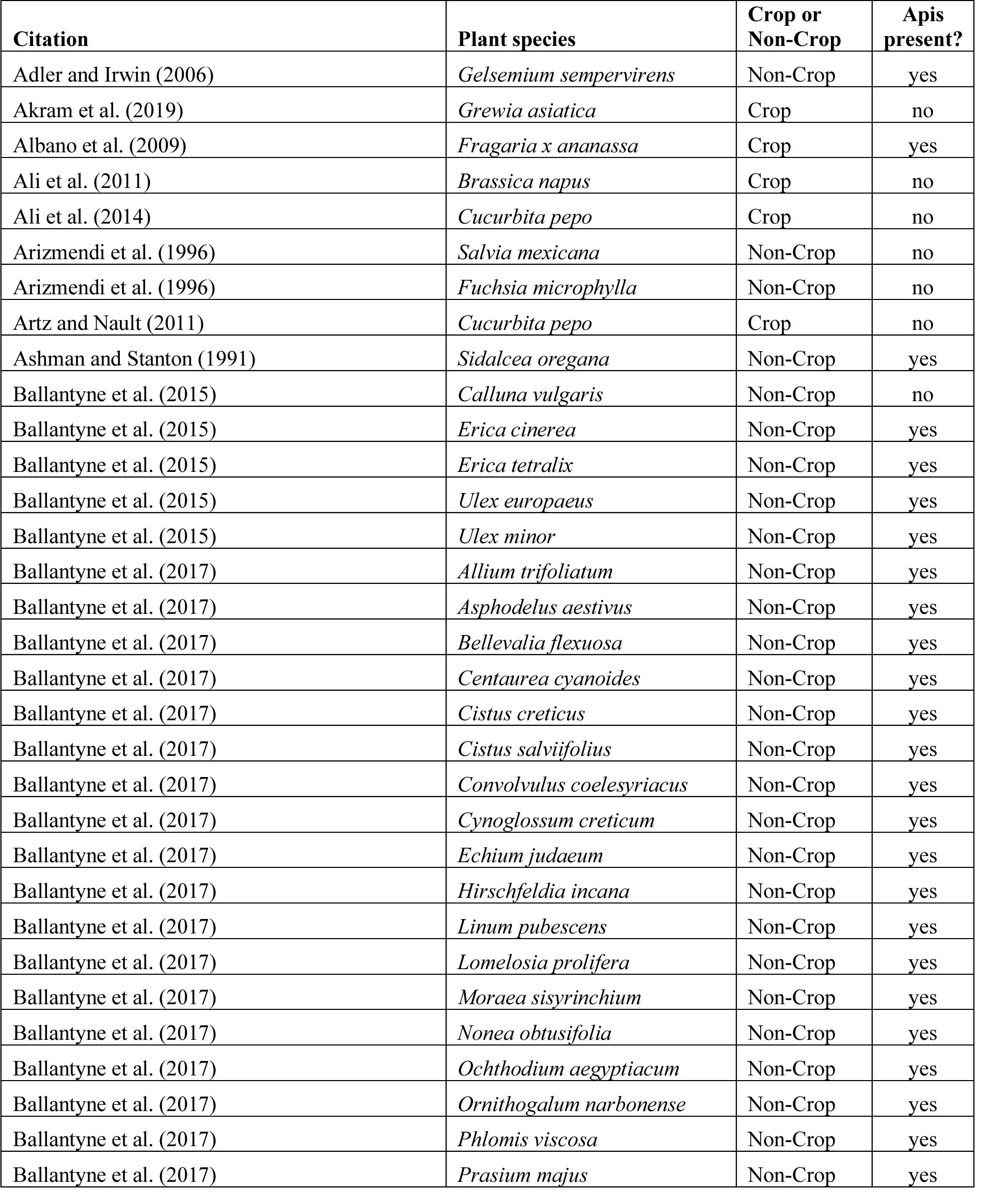
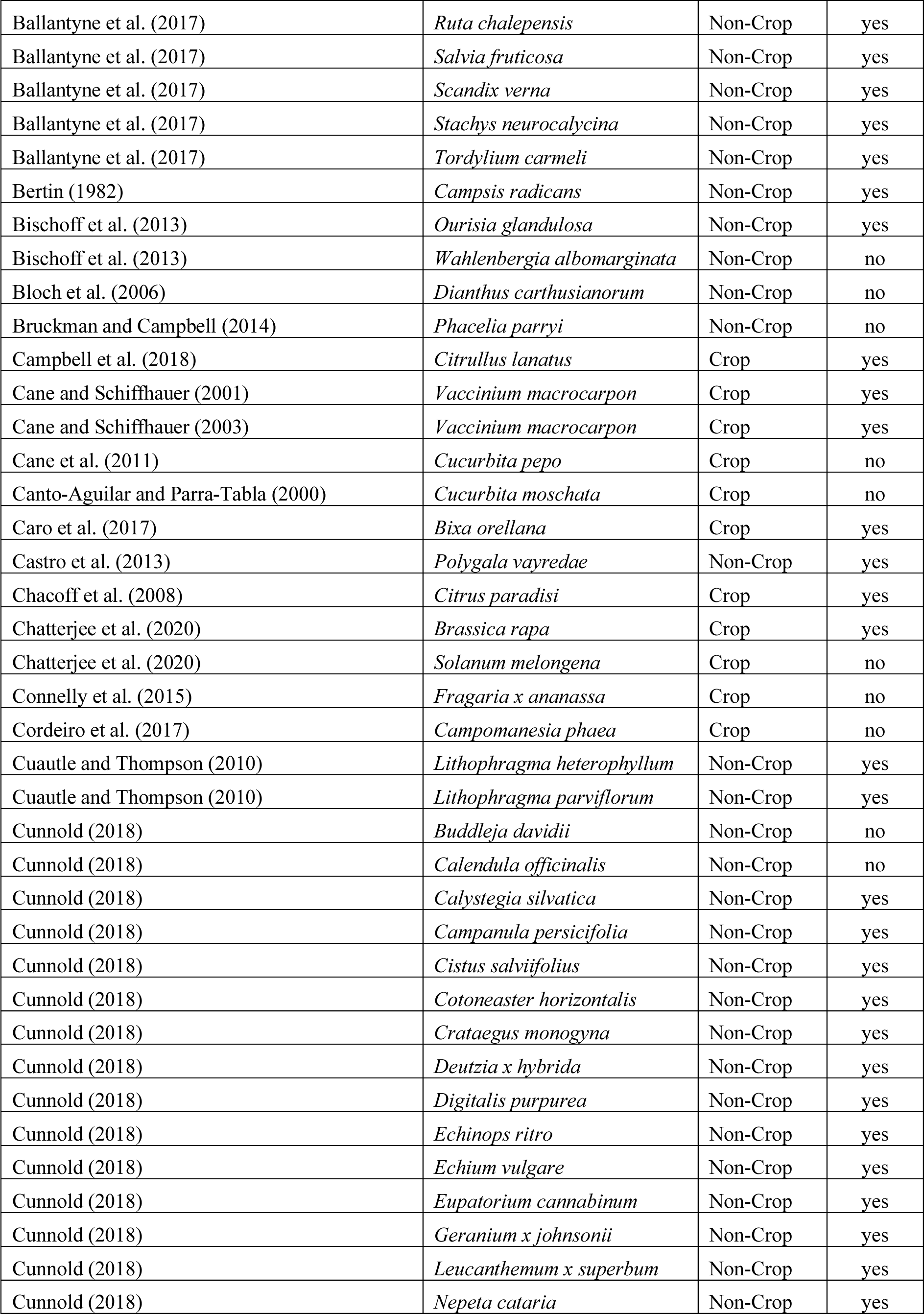
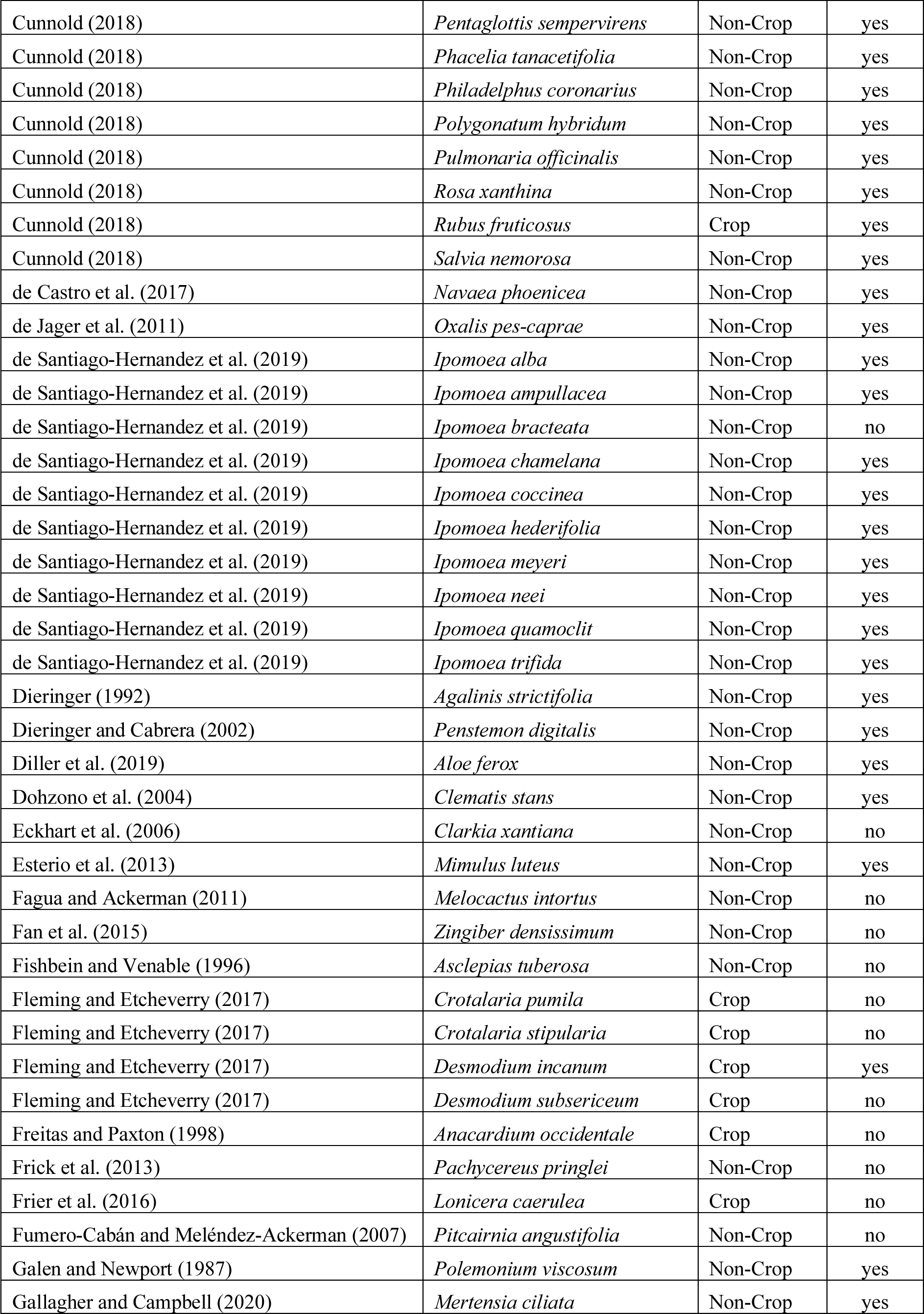
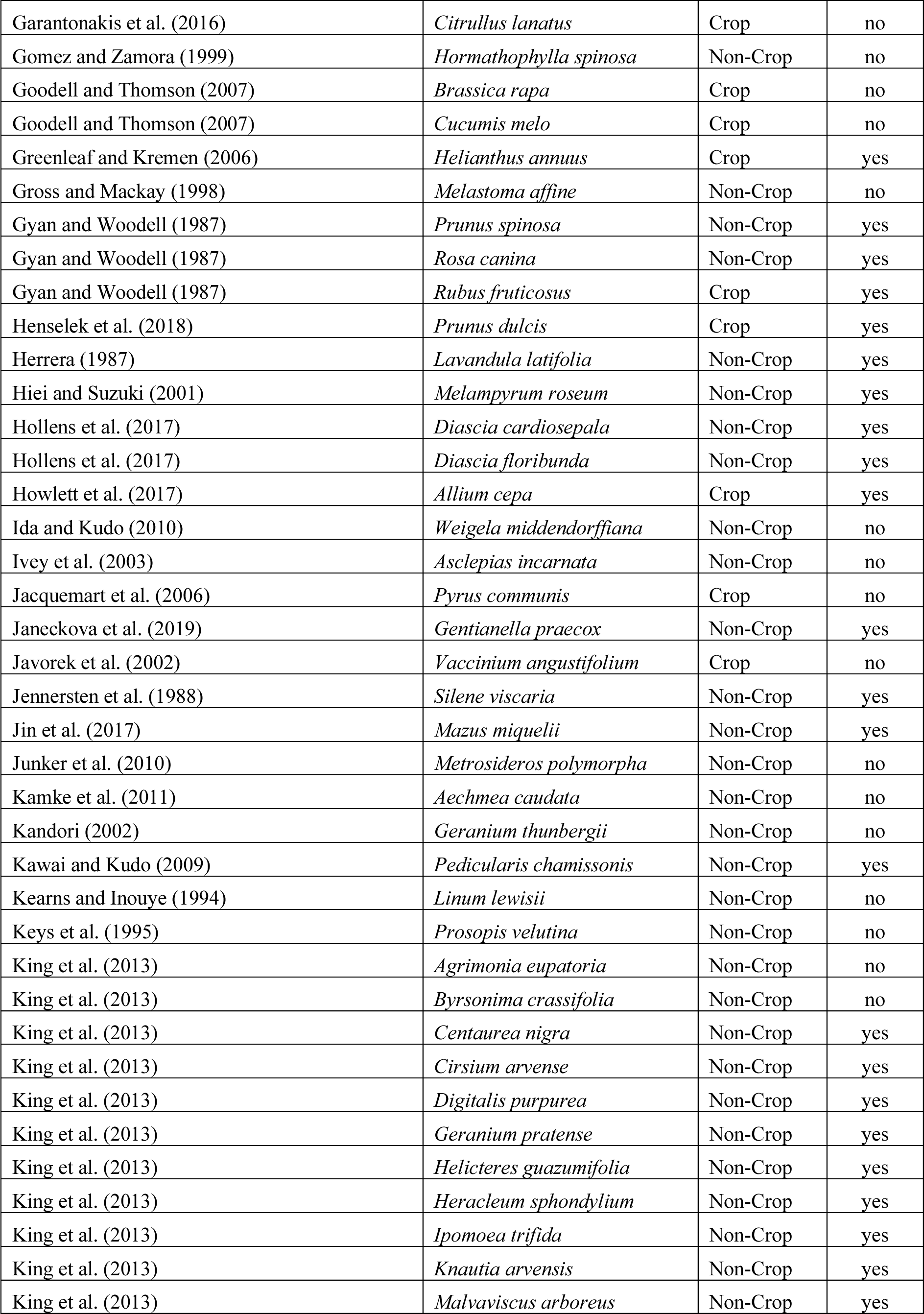
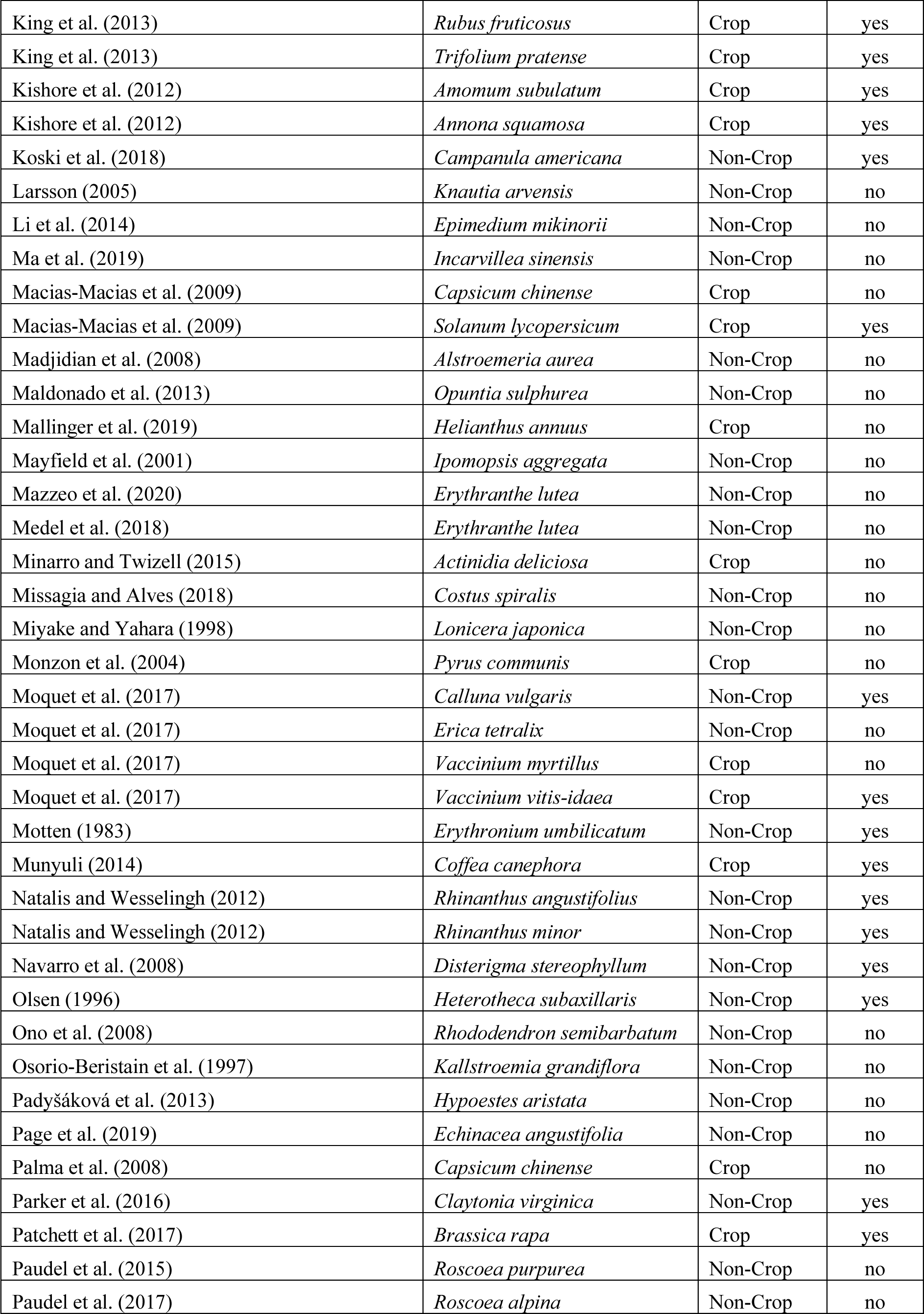
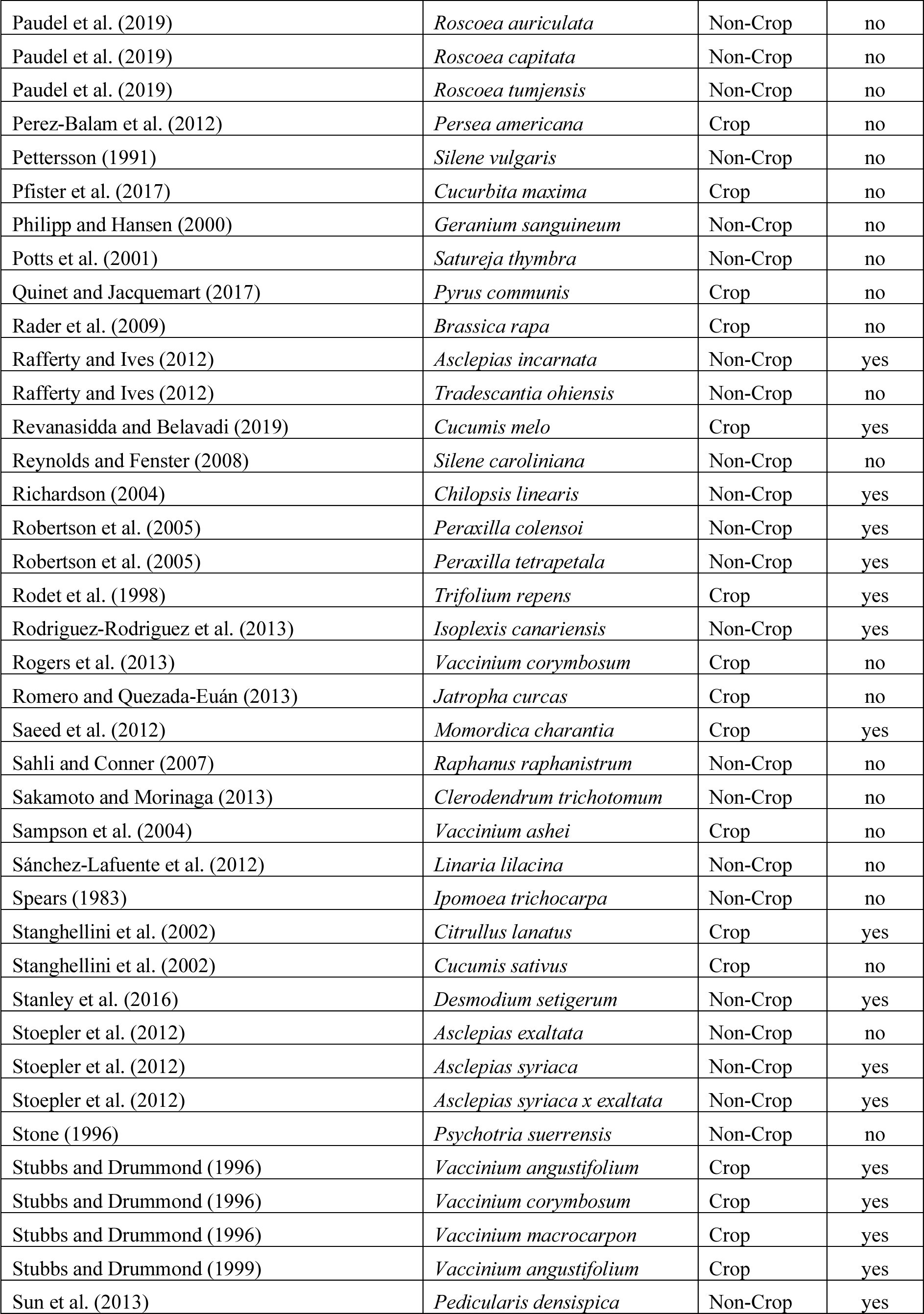
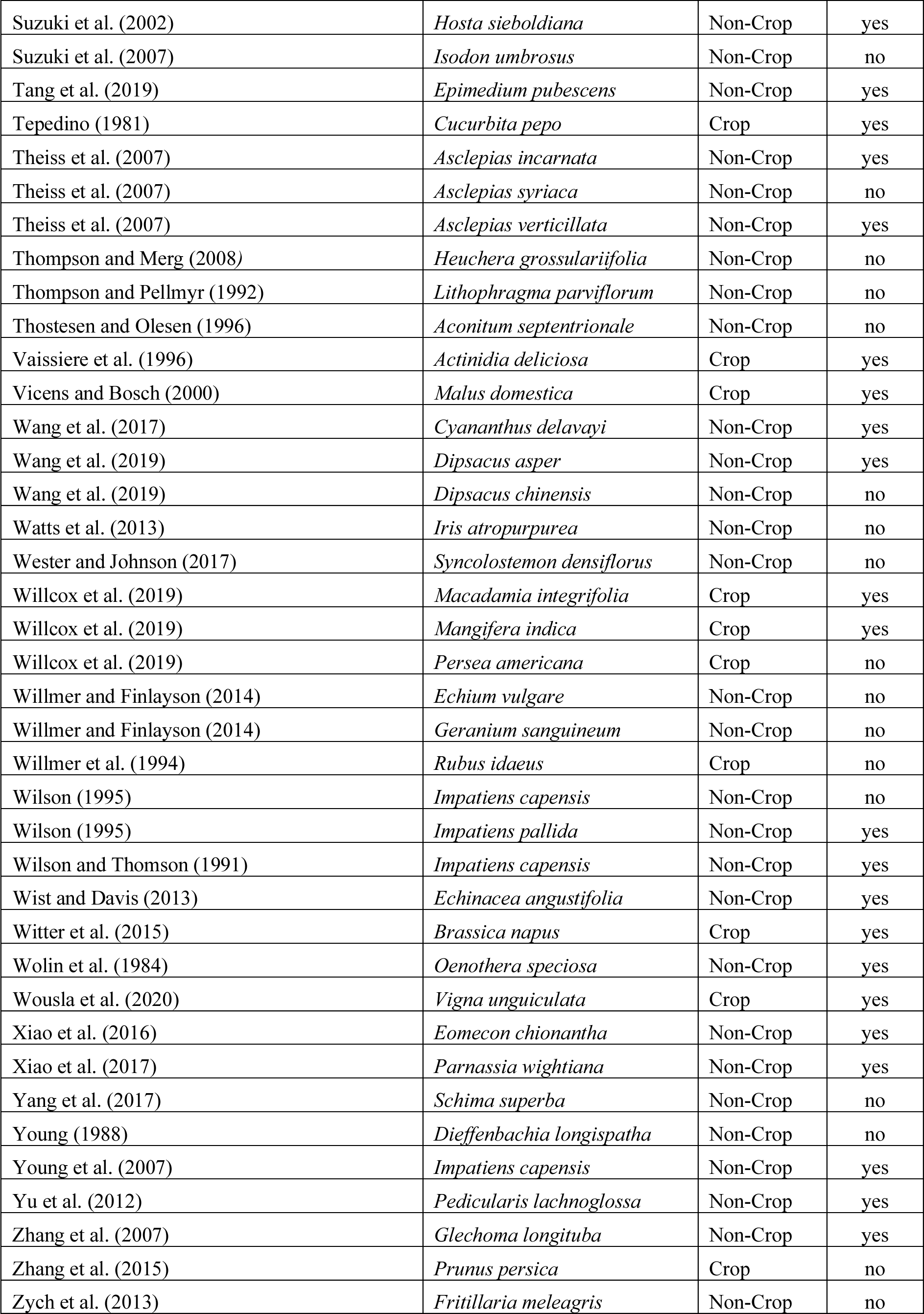
Studies included in the meta-analysis

**Figure S1.**
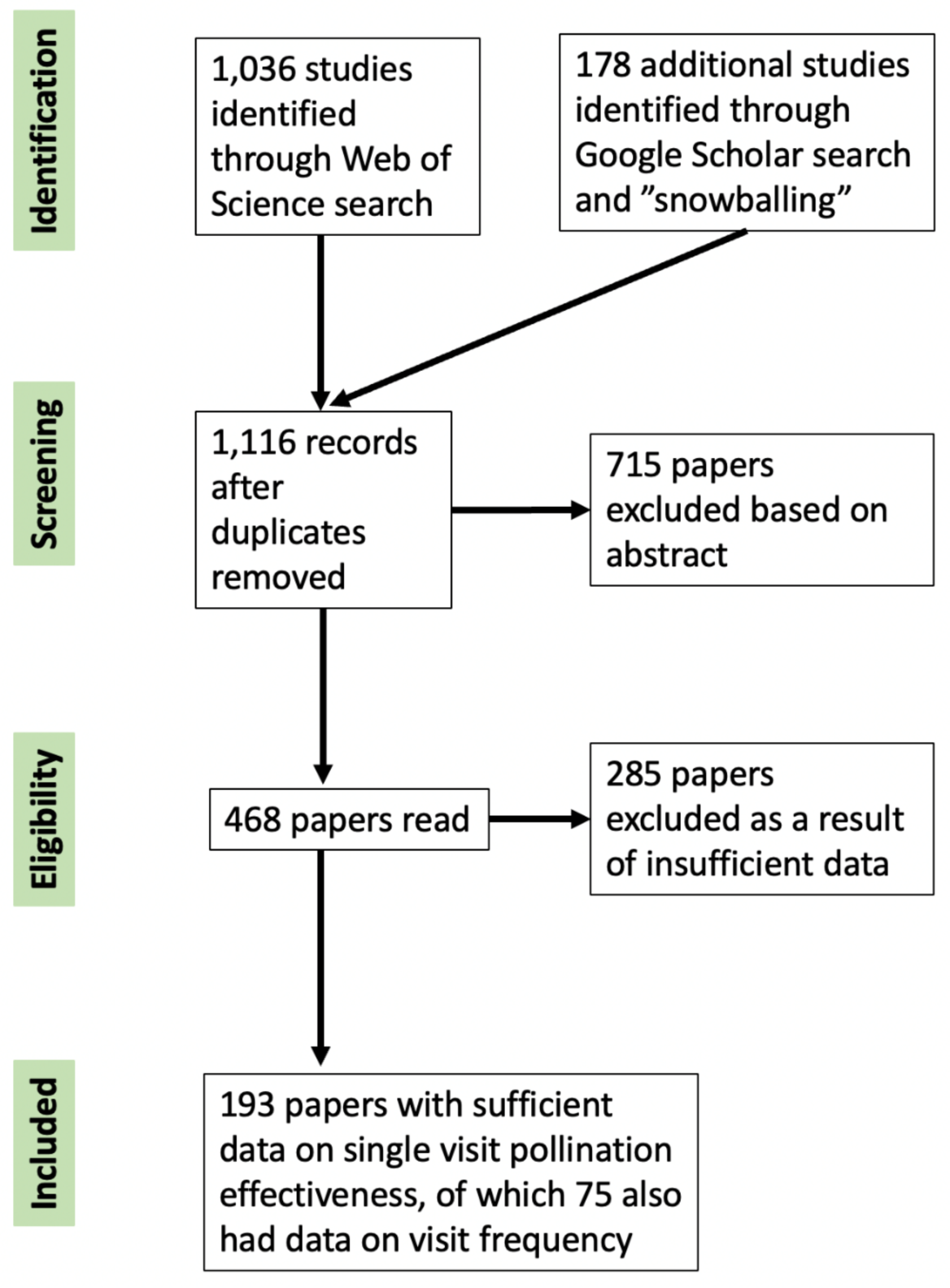
PRISMA diagram demonstrating the path through which papers were filtered for inclusion in the meta-analysis. We performed a Web of Science (WoS) search using the query: [“pollinat* effectiveness” OR “pollinat* efficacy” OR “pollinat* effectiveness” OR “pollinat* intensity” OR “pollinat* importance” OR “pollinat* level” OR “stigmatic fertilization success” “pollen transfer effect*” OR (“per visit” AND poll*) OR (“per-visit” AND poll*) OR (“per visit” AND seed) OR (“per-visit” AND seed) OR (“per visit” AND fruit) OR (“per-visit” AND fruit) OR (“single visit” AND fruit) OR (“single visit” AND seed) OR (“single visit” AND poll*)]. We performed a Google Scholar search using the keywords: (“single visit deposition”), (“per-visit” AND pollen), (pollinat* AND SVD), and (“pollen receipt” AND “per-visit”).

**Figure S2.**
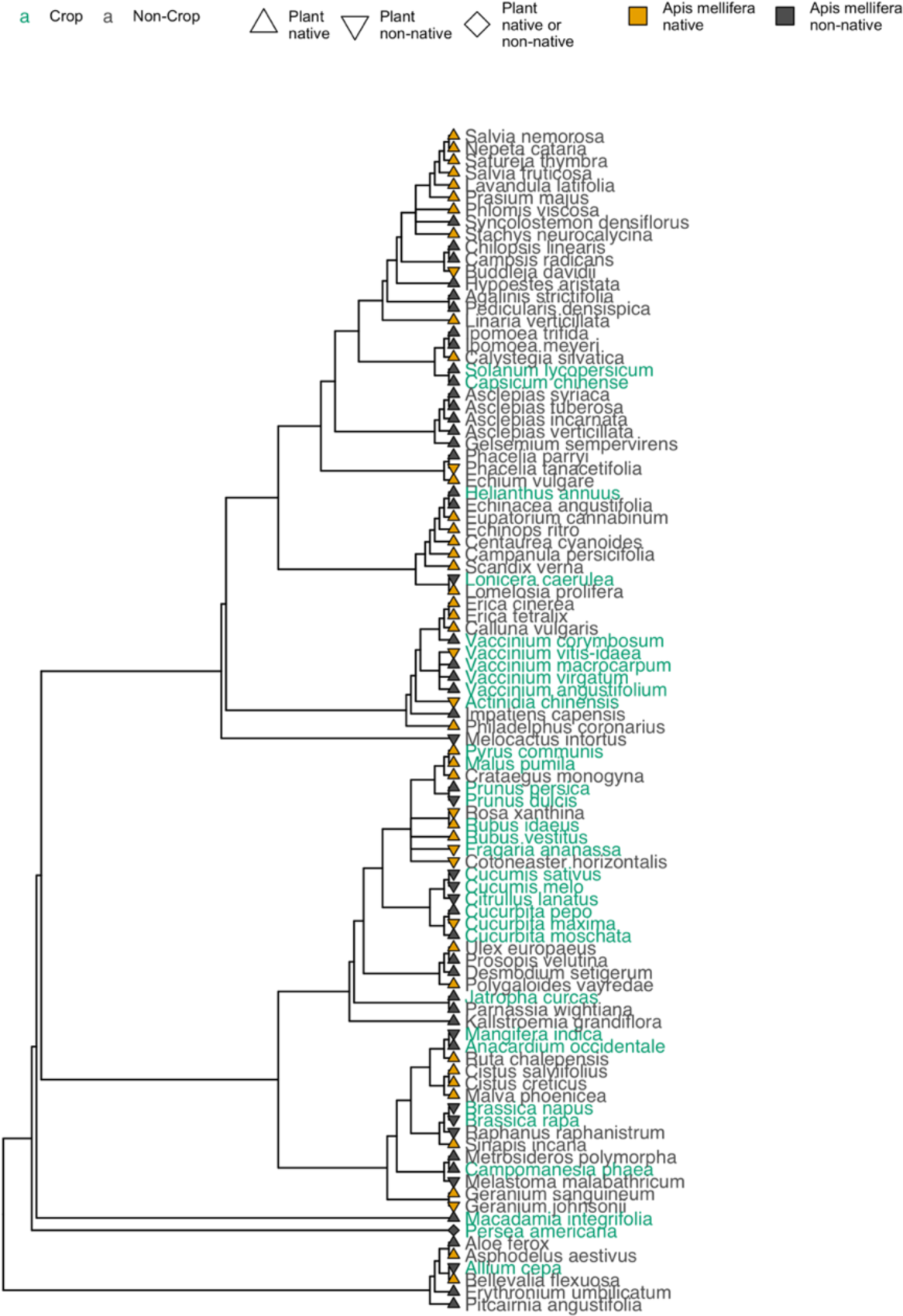
Studies with honeybee visitors explored single visit effectiveness in 252 plant species belonging to 67 families. Both crops (green text) and non-crops (black text) were examined outside (gray fill) and inside (orange fill) honeybees’ native range. These plant species were both native (triangles) and non-native (inverted triangles) to the regions in which they were studied. A few plant species were also investigated both inside and outside of their native range (diamonds). We included a phylogenetic covariance matrix based on this phylogeny as a random effect in all models.

**Figure S3.**
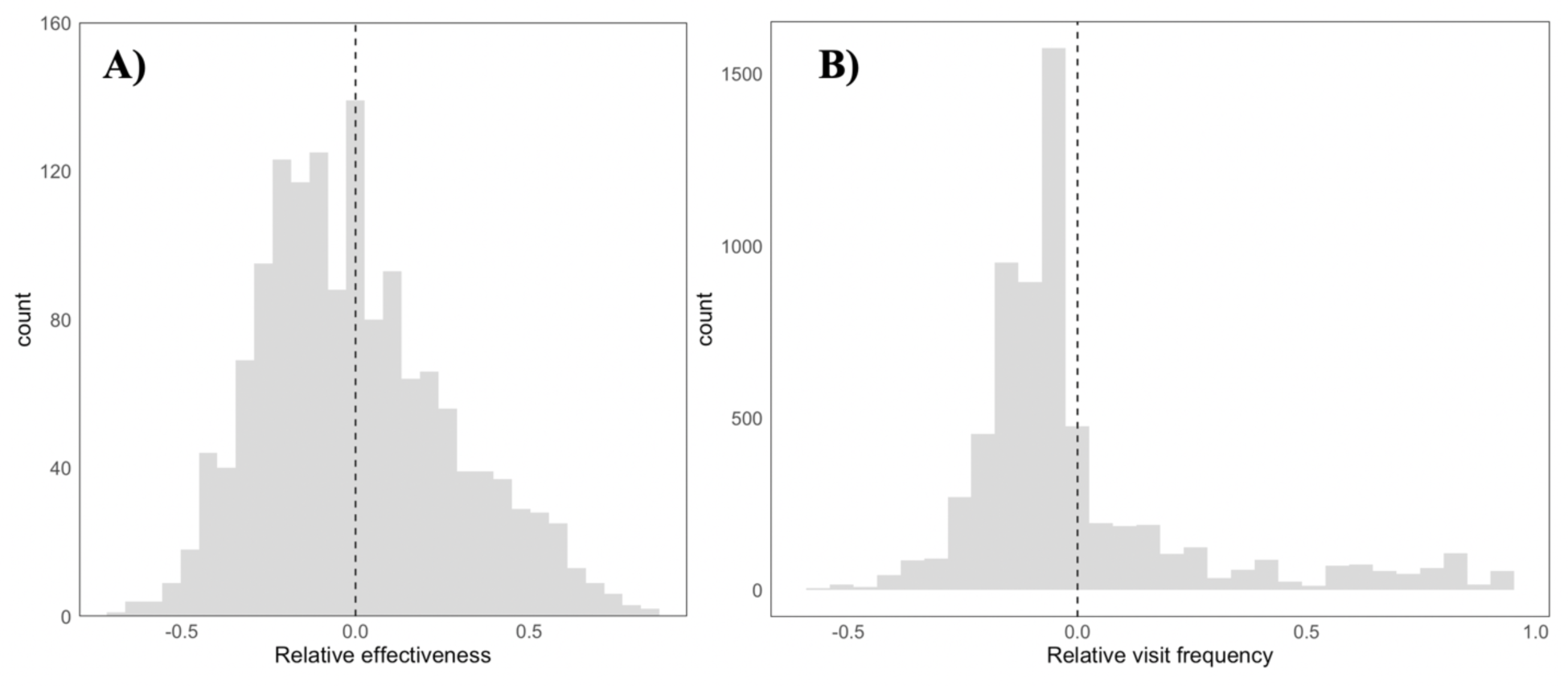
Histograms of (a) relative effectiveness values for all pollinators included in our meta-analysis and (b) the relative visit frequencies for all pollinators included in the subset of studies that reported paired data on visit frequencies and single visit effectiveness values. The relative effectiveness value is calculated as: (effectiveness value - mean effectiveness for unique study and plant)/maximum effectiveness for unique study and plant) such that positive values represent pollinators who were more effective that average and negative values represent pollinators who are less effective than average. Similarly, relative visit frequencies are calculated as: (visit value - visit value mean for unique study and plant)/maximum visit value for unique study and plant) such that positive values represent pollinators who visit more frequently than average and negative values represent pollinators who visit less frequently than average. Dividing by the maximum values for each unique study and plant ensures that the relativized effectiveness and visitation values are between -1 and 1 despite highly variable measures of visit frequency and effectiveness between studies.

**Figure S4.**
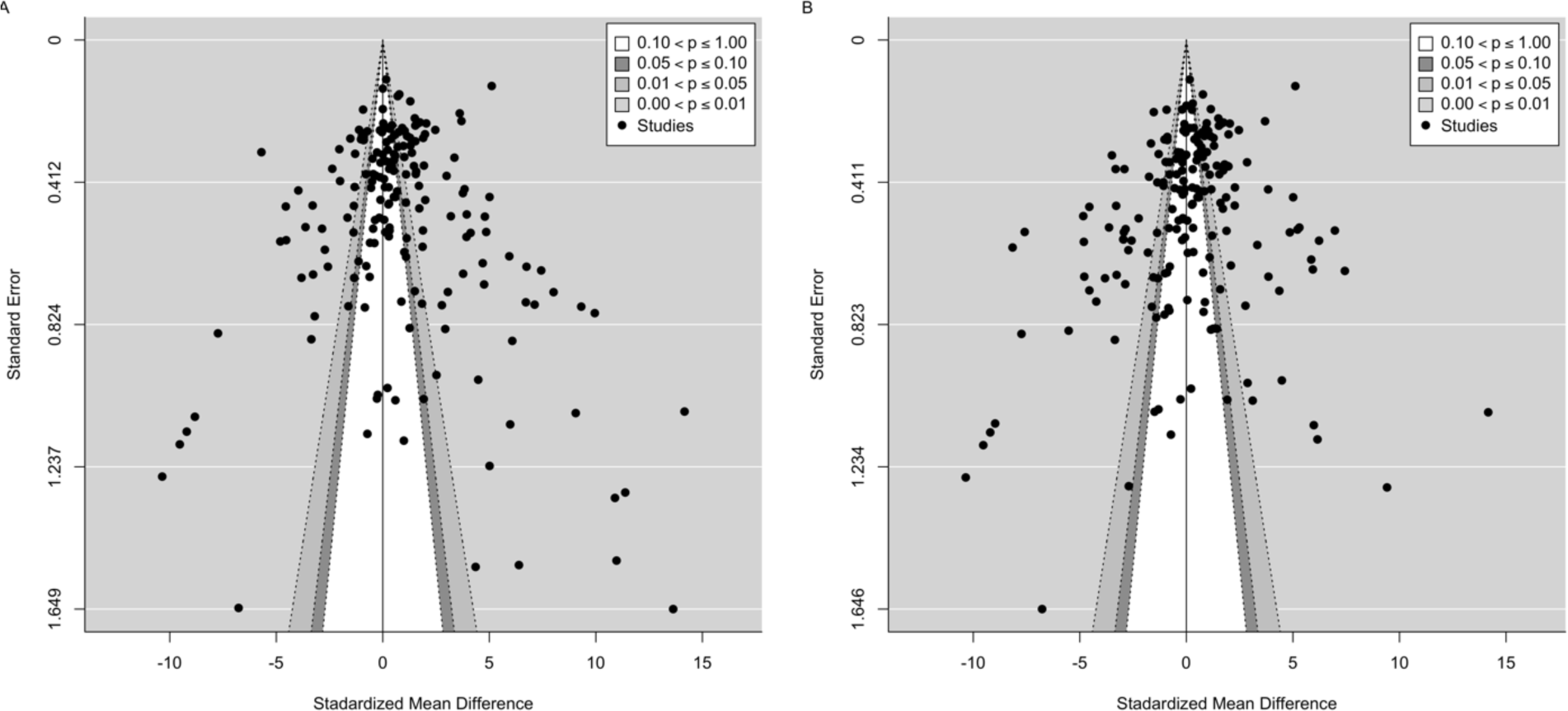
Funnel plots A) with most effective values and B) with most average values

**Figure S5.**
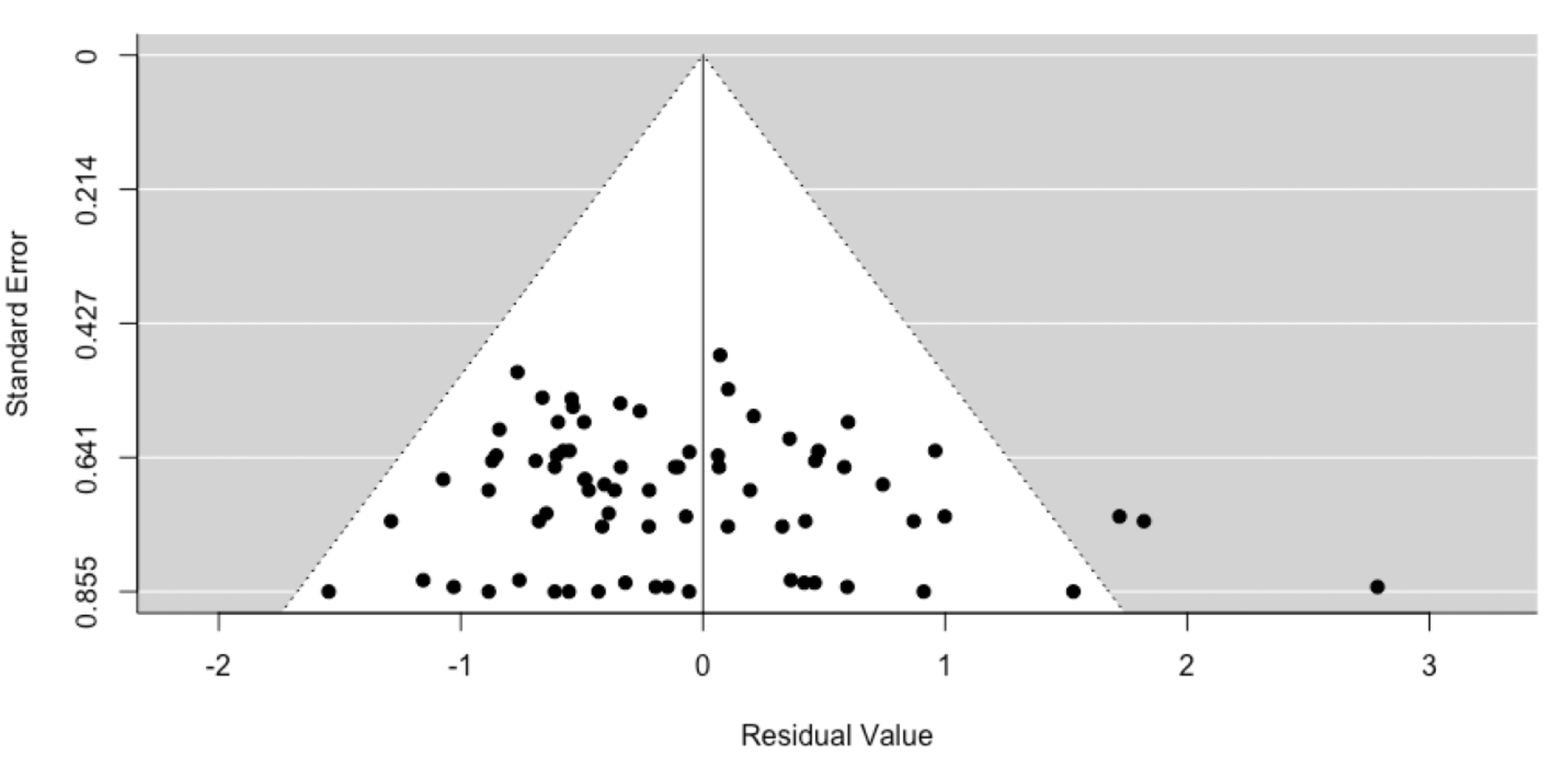
Funnel plot for the meta-regression comparing pollinator’s visit frequencies and single visit effectiveness.

**Figure S6.**
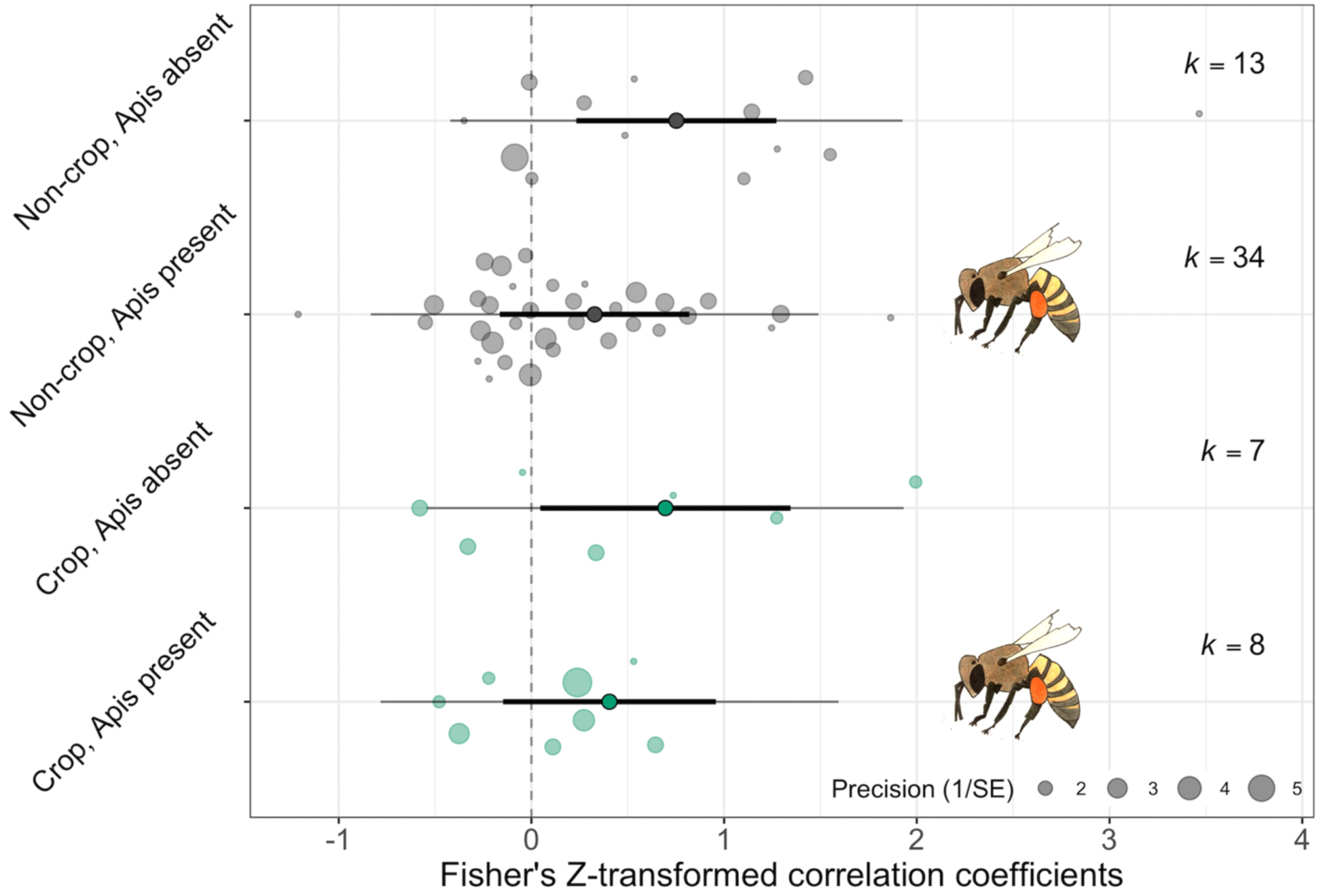
Meta-regression results for the relationship between a pollinator’s visit frequency and single visit effectiveness for crop and non-crop plants in studies with and without honeybees present. Effect sizes (Fisher’s Z-transformed correlation coefficients) were compared for non-crop (gray circles) and crop species (green circles) in studies where honeybees was present (as indicated by the honeybee icons) and systems where they were absent. Meta-analytic means are represented as point estimates with their 95% CI (thick lines) and prediction intervals (thin lines). Individual effect sizes are scaled by their precision (1/SE).

## Appendix S2

**Table S1.**
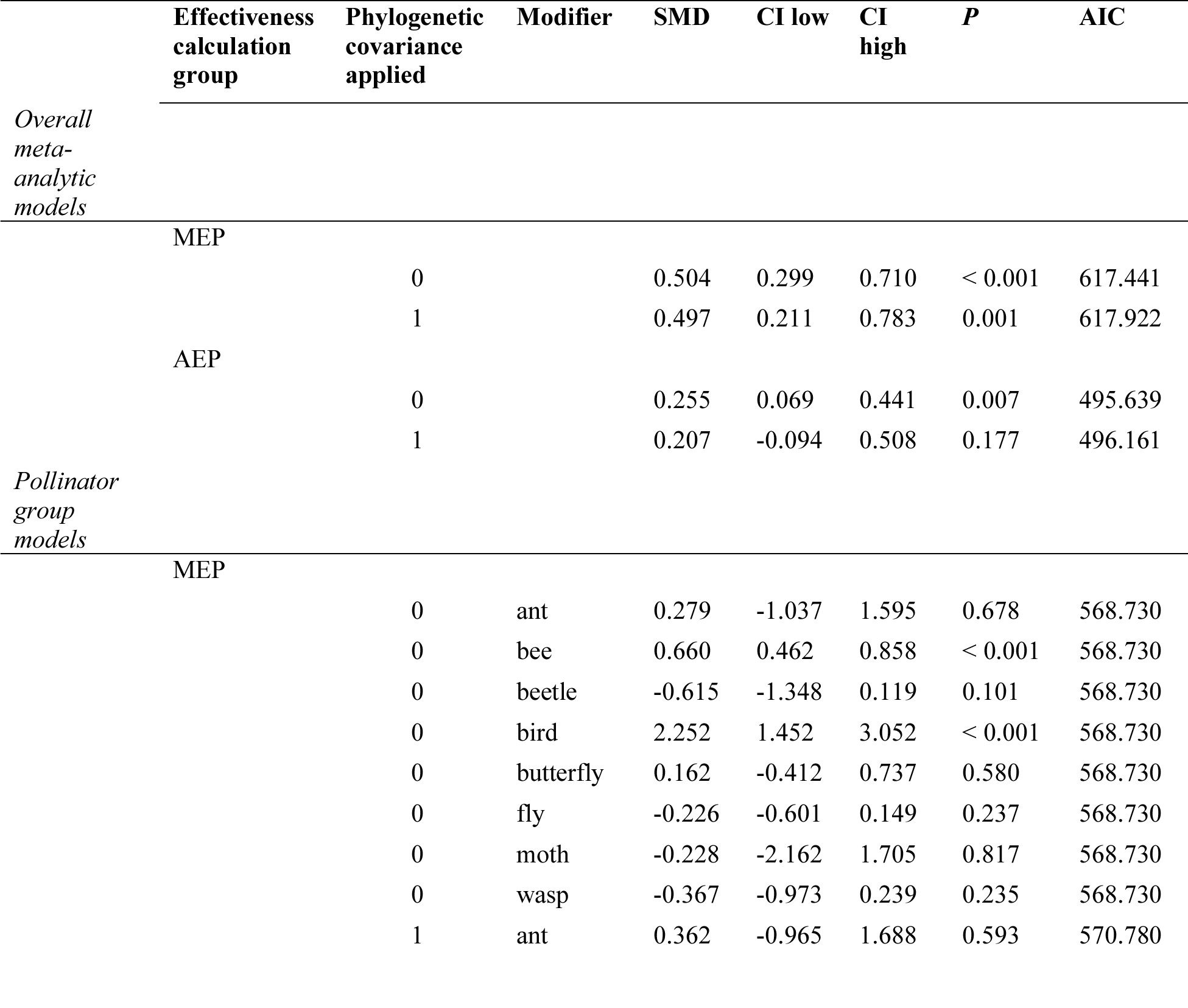
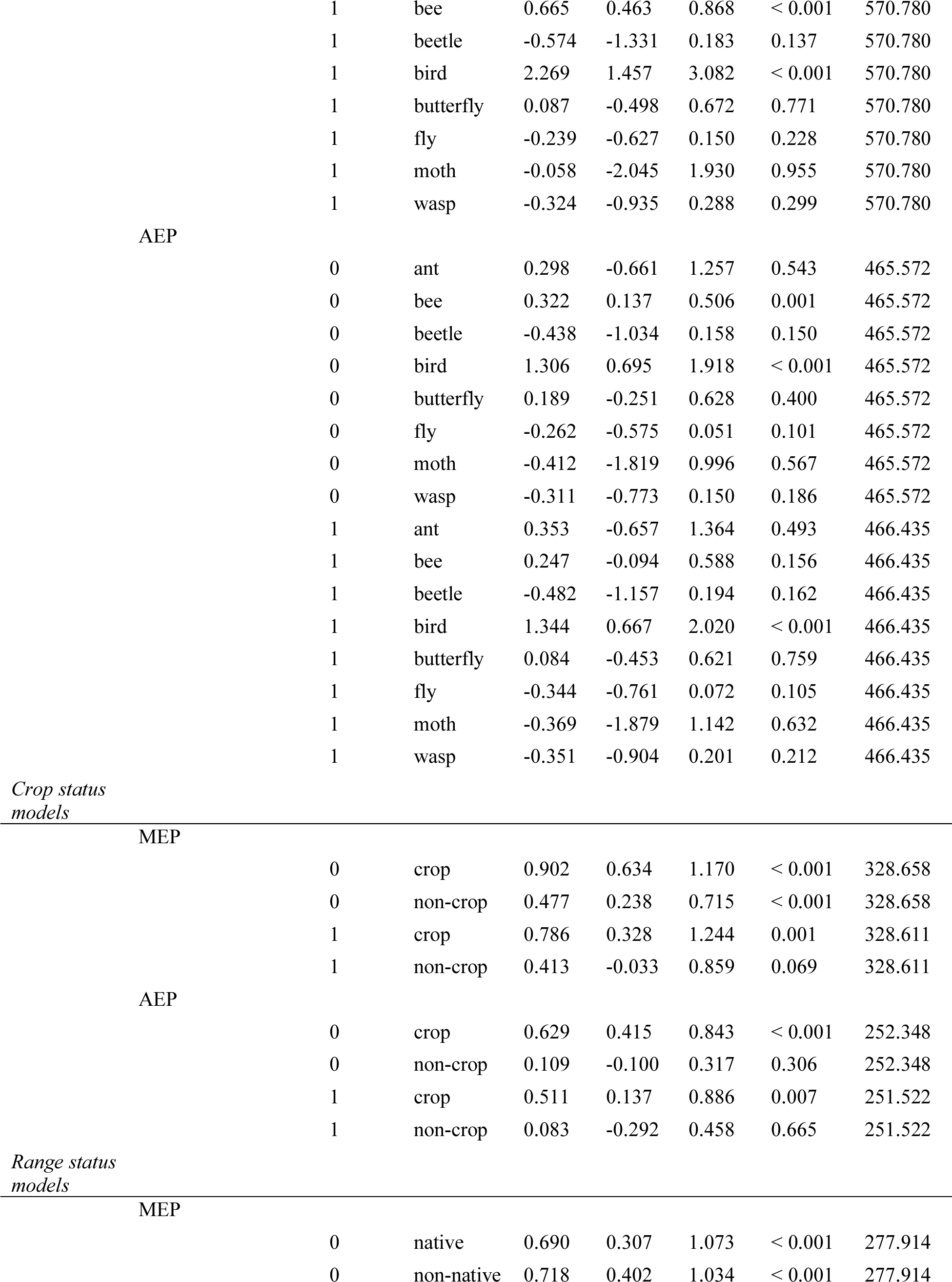
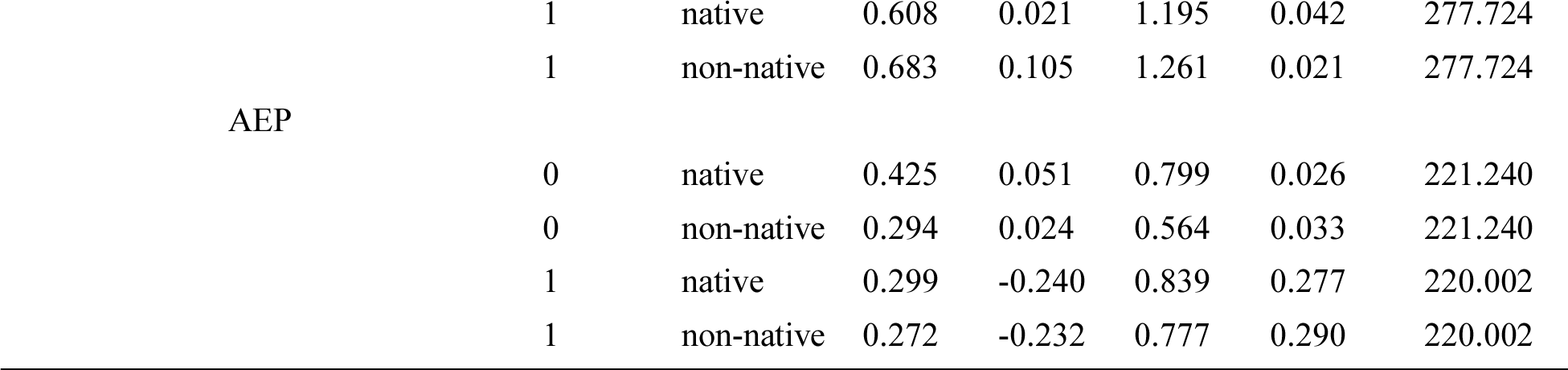
Model outputs for most effective (MEP) and average effectiveness (AEP) effect size calculations graphed in Fig. 2, 3, and 4. When phylogenetic covariance applied is ‘1’ this indicates that models included phylogenetic covariance matrices as random effects. When phylogenetic covariance applied is ‘0’ no such control was not included. All models had study ID, site, year, and plant species as random effects. Despite slightly higher AIC values and larger P values we present results from models including phylogenetic controls to fully account for non-independence due to shared ancestry.

## Literature Cited

1. Adler, L. S., and R. E. Irwin. 2006. Comparison of pollen transfer dynamics by multiple floral visitors: Experiments with pollen and fluorescent dye. Annals of Botany 97: 141–150.

2. Aizen, M. A., and L. D. Harder. 2009. The global stock of domesticated honeybees Is growing slower than agricultural demand for pollination. Current Biology 19: 915–918.

3. Ballantyne, G., K. C. R. Baldock, L. Rendell, and P. G. Willmer. 2017. Pollinator importance networks illustrate the crucial value of bees in a highly speciose plant community. Scientific Reports 7: 8389.

4. Ballantyne, G., K. C. R. Baldock, and P. G. Willmer. 2015. Constructing more informative plant-pollinator networks: visitation and pollen deposition networks in a heathland plant community. Proceedings of the Royal Society B-Biological Sciences 282: 14–22.

5. Blitzer, E. J., J. Gibbs, M. G. Park, and B. N. Danforth. 2016. Pollination services for apple are dependent on diverse wild bee communities. Agriculture, Ecosystems & Environment 221: 1–7.

6. Bosch, J., W. P. Kemp, and G. E. Trostle. 2006. Bee population returns and cherry yields in an orchard pollinated with *Osmia lignaria* (Hymenoptera: Megachilidae). Journal of Economic Entomology 99: 408–413.

7. Brosi, B. J., and H. M. Briggs. 2013. Single pollinator species losses reduce floral fidelity and plant reproductive function. Proceedings of the National Academy of Sciences 110: 13044–13048.

8. Cane, J. H., and V. J. Tepedino. 2017. Gauging the effect of honeybee pollen collection on native bee communities. Conservation Letters 10: 205–210.

9. Cardinale, B. J., J. E. Duffy, A. Gonzalez, D. U. Hooper, C. Perrings, P. Venail, A. Narwani, G. M. Mace, D. Tilman, D. A. Wardle, A. P. Kinzig, G. C. Daily, M. Loreau, J. B. Grace, A. Larigauderie, D. S. Srivastava, and S. Naeem. 2012. Biodiversity loss and its impact on humanity. Nature 486: 59–67.

10. Castellanos, M. C., P. Wilson, S. J. Keller, A. D. Wolfe, and J. D. Thomson. 2006. Anther evolution: pollen presentation strategies when pollinators differ. The American Naturalist 167: 288–296.

11. Chamberlain, S. A., S. M. Hovick, C. J. Dibble, N. L. Rasmussen, B. G. V. Allen, B. S. Maitner, J. R. Ahern, L. P. Bell-Dereske, C. L. Roy, M. Meza-Lopez, J. Carrillo, E. Siemann, M. J. Lajeunesse, and K. D. Whitney. 2012. Does phylogeny matter? Assessing the impact of phylogenetic information in ecological meta-analysis. Ecology Letters 15: 627–636.

12. Chamberlain, S. 2020. brranching: Fetch “Phylogenies” from many sources, version 0.6.0. https://CRAN.R-project.org/package=brranching.

13. Cornwell, W., and S. Nakagawa. 2017. Phylogenetic comparative methods. Current Biology 27: R333–R336.

14. Egger, M., G. D. Smith, M. Schneider, and C. Minder. 1997. Bias in meta-analysis detected by a simple, graphical test. BMJ 315: 629–634.

15. Földesi, R., B. G. Howlett, I. Grass, and P. Batáry. 2020. Larger pollinators deposit more pollen on stigmas across multiple plant species—A meta-analysis. Journal of Applied Ecology 00: 1–9.

16. Fontaine, C., I. Dajoz, J. Meriguet, and M. Loreau. 2005. Functional diversity of plant–pollinator interaction webs enhances the persistence of plant communities. PLOS Biology 4: e1.

17. Gómez, J. M., M. Abdelaziz, J. Lorite, A. J. Muñoz-Pajares, and F. Perfectti. 2010. Changes in pollinator fauna cause spatial variation in pollen limitation. Journal of Ecology 98: 1243– 1252.

18. Goodell, K., and J. D. Thomson. 1997. Comparisons of pollen removal and deposition by honeybees and bumblebees visiting apple. Acta Horticulturae 437: 103–108.

19. Grafen, A., and W. D. Hamilton. 1989. The phylogenetic regression. Philosophical Transactions of the Royal Society of London. B, Biological Sciences 326: 119–157.

20. Hansen, D. M., J. M. Olesen, and C. G. Jones. 2002. Trees, birds and bees in Mauritius: exploitative competition between introduced honey bees and endemic nectarivorous birds? Journal of Biogeography 29: 721–734.

21. Harder, L. D., and S. C. H. Barrett. 1995. Mating cost of large floral displays in hermaphrodite plants. Nature 373: 512–515.

22. Harder, L. D., and J. D. Thomson. 1989. Evolutionary options for maximizing pollen dispersal of animal-pollinated plants. The American Naturalist 133: 323–344.

23. Harrison, T., J. Gibbs, and R. Winfree. 2018. Forest bees are replaced in agricultural and urban landscapes by native species with different phenologies and life-history traits. Global Change Biology 24: 287–296.

24. Hedges, L. V. 1981. Distribution theory for Glass’s estimator of effect size and related estimators. Journal of Educational Statistics 6: 107–128.

25. Herrera, C. M. 1987. Components of pollinator “quality”: comparative analysis of a diverse insect assemblage. Oikos 50: 79–90.

26. Herrera, C. M. 2020. Gradual replacement of wild bees by honeybees in flowers of the Mediterranean Basin over the last 50 years. Proceedings of the Royal Society B: Biological Sciences 287: 20192657.

27. Hung, K.-L. J., J. M. Kingston, M. Albrecht, D. A. Holway, and J. R. Kohn. 2018. The worldwide importance of honeybees as pollinators in natural habitats. Proceedings of the Royal Society B: Biological Sciences 285: 20172140.

28. Inouye, D. W., D. E. Gill, M. R. Dudash, and C. B. Fenster. 1994. A model and lexicon for pollen fate. American Journal of Botany 81: 1517–1530.

29. Irwin, R. E., J. L. Bronstein, J. S. Manson, and L. Richardson. 2010. Nectar robbing: ecological and evolutionary perspectives. Annual Review of Ecology, Evolution, and Systematics 41: 271–292.

30. Isaacs, R., N. Williams, J. Ellis, T. L. Pitts-Singer, R. Bommarco, and M. Vaughan. 2017. Integrated crop pollination: combining strategies to ensure stable and sustainable yields of pollination-dependent crops. Basic and Applied Ecology 22: 44–60.

31. Javorek, S. K., K. E. Mackenzie, and S. P. V. Kloet. 2002. Comparative pollination effectiveness among bees (Hymenoptera: Apoidea) on Lowbush blueberry (Ericaceae: *Vaccinium angustifolium*). Annals of the Entomological Society of America 95: 345–351.

32. King, C., G. Ballantyne, and P. G. Willmer. 2013. Why flower visitation is a poor proxy for pollination: measuring single-visit pollen deposition, with implications for pollination networks and conservation. Methods in Ecology and Evolution 4: 811–818.

33. Koricheva, J., J. Gurevitch, and K. Mengersen. 2013. Handbook of Meta-analysis in Ecology and Evolution. Princeton University Press.

34. Lohbeck, M., F. Bongers, M. Martinez-Ramos, and L. Poorter. 2016. The importance of biodiversity and dominance for multiple ecosystem functions in a human-modified tropical landscape. Ecology 97: 2772–2779.

35. Loreau, M., S. Naeem, P. Inchausti, J. Bengtsson, J. P. Grime, A. Hector, D. U. Hooper, M. A. Huston, D. Raffaelli, B. Schmid, D. Tilman, and D. A. Wardle. 2001. Biodiversity and ecosystem functioning: current knowledge and future challenges. Science 294: 804–808.

36. Mallinger, R. E., and C. Gratton. 2015. Species richness of wild bees, but not the use of managed honeybees, increases fruit set of a pollinator-dependent crop. Journal of Applied Ecology 52: 323–330.

37. Minnaar, C., B. Anderson, M. L. de Jager, and J. D. Karron. 2019. Plant–pollinator interactions along the pathway to paternity. Annals of Botany 123: 225–245.

38. Moher, D., A. Liberati, J. Tetzlaff, D. G. Altman, and T. P. Group. 2009. Preferred reporting items for systematic reviews and meta-analyses: The PRISMA statement. PLOS Medicine 6: e1000097.

39. Morales, C. L., and A. Traveset. 2008. Interspecific pollen transfer: magnitude, prevalence and consequences for plant Fitness. Critical Reviews in Plant Sciences 27: 221–238.

40. Nakagawa, S., and E. S. A. Santos. 2012. Methodological issues and advances in biological meta-analysis. Evolutionary Ecology 26: 1253–1274.

41. Ne’eman, G., A. Jürgens, L. Newstrom-Lloyd, S. G. Potts, and A. Dafni. 2010. A framework for comparing pollinator performance: effectiveness and efficiency. Biological Reviews 85: 435–451.

42. Ohara, M., S. Higashi, and A. Ohara. 1994. Effects of inflorescence size on visits from pollinators and seed set of *Corydalis ambigua* (Papaveraceae). Oecologia 98: 25–30.

43. Ollerton, J., R. Winfree, and S. Tarrant. 2011. How many flowering plants are pollinated by animals? Oikos 120: 321–326.

44. Page, M. L., J. L. Ison, A. L. Bewley, K. M. Holsinger, A. D. Kaul, K. E. Koch, K. M. Kolis, and S. Wagenius. 2019. Pollinator effectiveness in a composite: a specialist bee pollinates more florets but does not move pollen farther than other visitors. American Journal of Botany 106: 1487–1498.

45. Paradis, E., and K. Schliep. 2019. ape 5.0: an environment for modern phylogenetics and evolutionary analyses in R. Bioinformatics 35: 526–528.

46. Parker, A. J., N. M. Williams, and J. D. Thomson. 2016. Specialist pollinators deplete pollen in the spring ephemeral wildflower *Claytonia virginica*. Ecology and Evolution 6: 5169– 5177.

47. Pérez-Méndez, N., G. K. S. Andersson, F. Requier, J. Hipólito, M. A. Aizen, C. L. Morales, N. García, G. P. Gennari, and L. A. Garibaldi. 2020. The economic cost of losing native pollinator species for orchard production. Journal of Applied Ecology 57: 599–608.

48. R Core Team. 2020. R: A language and environment for statistical computing. R Foundation for Statistical Computing, Vienna, Austria. https://www.R-project.org/.

49. Rader, R., J. Reilly, I. Bartomeus, and R. Winfree. 2013. Native bees buffer the negative impact of climate warming on honey bee pollination of watermelon crops. Global Change Biology 19: 3103–3110.

50. Rader, R., S. A. Cunningham, B. G. Howlett, and D. W. Inouye. 2020. Non-bee insects as visitors and pollinators of crops: Biology, ecology, and management. Annual Review of Entomology 65: 391–407.

51. Rohatgi, A. 2020. WebPlotDigitizer, Version 4.4. https://automeris.io/WebPlotDigitizer.

52. Ruttner, F. 1988. Biogeography and taxonomy of honeybees. Berlin, Germany. Springer.

53. Sáez, A., C. L. Morales, L. Y. Ramos, and M. A. Aizen. 2014. Extremely frequent bee visits increase pollen deposition but reduce drupelet set in raspberry. Journal of Applied Ecology 51: 1603–1612.

54. Sterne, J. A. C., and M. Egger. 2005. Regression Methods to Detect Publication and Other Bias in Meta-Analysis. Pages 99–110 in H.R. Rothstein, A.J. Sutton, and M. Borenstein, editors. Publication Bias in Meta-Analysis: Prevention Assessment and Adjustments. John Wiley & Sons, Ltd.

55. Sun, S.-G., S.-Q. Huang, and Y.-H. Guo. 2013. Pollinator shift to managed honeybees enhances reproductive output in a bumblebee-pollinated plant. Plant Systematics and Evolution 299: 139–150.

56. Thomson, J. D. 1986. Pollen transport and deposition by bumble bees in Erythronium: Influences of floral nectar and bee grooming. Journal of Ecology 74: 329–341.

57. Valido, A., M. C. Rodríguez-Rodríguez, and P. Jordano. 2019. Honeybees disrupt the structure and functionality of plant-pollinator networks. Scientific Reports 9: 4711.

58. Vázquez, D. P., S. B. Lomáscolo, M. B. Maldonado, N. P. Chacoff, J. Dorado, E. L. Stevani, and N. L. Vitale. 2012. The strength of plant–pollinator interactions. Ecology 93: 719–725.

59. Vicens, N., and J. Bosch. 2000. Pollinating efficacy of *Osmia cornuta* and *Apis mellifera* (Hymenoptera: Megachilidae, Apidae) on ‘Red Delicious’ apple. Environmental Entomology 29: 235–240.

60. Viechtbauer, W. 2010. Conducting meta-analyses in R with the metafor package. Journal of Statistical Software 36: 1–48.

61. Westerkamp, C. 1991. Honeybees are poor pollinators — why? Plant Systematics and Evolution 177: 71–75.

62. Willcox, B. K., M. A. Aizen, S. A. Cunningham, M. M. Mayfield, and R. Rader. 2017. Deconstructing pollinator community effectiveness. Current Opinion in Insect Science 21: 98–104.

63. Wilson, P., and J. D. Thomson. 1991. Heterogeneity among floral visitors leads to discordance between removal and deposition of pollen. Ecology 72: 1503–1507.

64. Winfree, R., J. R. Reilly, I. Bartomeus, D. P. Cariveau, N. M. Williams, and J. Gibbs. 2018. Species turnover promotes the importance of bee diversity for crop pollination at regional scales. Science 359: 791–793.

65. Winfree, R., J. W. Fox, N. M. Williams, J. R. Reilly, and D. P. Cariveau. 2015. Abundance of common species, not species richness, drives delivery of a real-world ecosystem service. Ecology Letters 18: 626–635.

66. Wist, T. J., and A. R. Davis. 2013. Evaluation of inflorescence visitors as pollinators of *Echinacea angustifolia* (Asteraceae): comparison of techniques. Journal of Economic Entomology 106: 2055–2071.

67. Woodcock, B. A., M. P. D. Garratt, G. D. Powney, R. F. Shaw, J. L. Osborne, J. Soroka, S. a. M. Lindström, D. Stanley, P. Ouvrard, M. E. Edwards, F. Jauker, M. E. McCracken, Y. Zou, S. G. Potts, M. Rundlöf, J. A. Noriega, A. Greenop, H. G. Smith, R. Bommarco, W. van der Werf, J. C. Stout, I. Steffan-Dewenter, L. Morandin, J. M. Bullock, and R. F. Pywell. 2019. Meta-analysis reveals that pollinator functional diversity and abundance enhance crop pollination and yield. Nature Communications 10: 1481.

68. Zanne, A. E., D. C. Tank, W. K. Cornwell, J. M. Eastman, S. A. Smith, R. G. FitzJohn, D. J. McGlinn, B. C. O’Meara, A. T. Moles, P. B. Reich, D. L. Royer, D. E. Soltis, P. F. Stevens, M. Westoby, I. J. Wright, L. Aarssen, R. I. Bertin, A. Calaminus, R. Govaerts, F. Hemmings, M. R. Leishman, J. Oleksyn, P. S. Soltis, N. G. Swenson, L. Warman, and J. M. Beaulieu. 2014. Three keys to the radiation of angiosperms into freezing environments. Nature 506: 89–92.

69. Zhang, H., J. Huang, P. H. Williams, B. E. Vaissière, Z. Zhou, Q. Gai, J. Dong, and J. An. 2015.Managed bumblebees outperform honeybees in increasing peach fruit set in China: different limiting processes with different pollinators. PLOS ONE 10: e0121143.

